# Functional connectome linking child-parent relationships with psychological problems in adolescence

**DOI:** 10.1101/678714

**Authors:** Takashi Itahashi, Naohiro Okada, Shuntaro Ando, Syudo Yamasaki, Daisuke Koshiyama, Kentaro Morita, Noriaki Yahata, Shinsuke Koike, Atsushi Nishida, Kiyoto Kasai, Ryu-ichiro Hashimoto

## Abstract

Paternal- and maternal-child relationships are associated with partly distinct psychobehavioral problems, which often manifest differently between boys and girls. In order to understand neural mechanisms underlying complicated mappings between child-parent relationship and adolescents’ problems, we used a dataset of early adolescents (*N*=93) and separately estimated the effects of paternal- and maternal-child relationships on resting-state functional connectivity and problems in boys and girls. General linear models identified the effects of paternal- and maternal-child relationships in different sets of functional connectivity, which we termed functional brain connectomes associated with paternal- and maternal-child relationship (FBCp and FBCm), respectively. Subsequent connectome-based models using these FBCs significantly predicted aggressive behaviors only for boys and internalizing problems selectively only girls. Lastly, a causal discovery method identified causal paths from daughter-mother relationship to FBCm, and then to daughter’s internalizing problems. These observations highlight sex-dependent mechanisms linking child-parent relationship, brain, and psychobehavioral problems in early adolescence.

---

Adolescence is a critical transition period from childhood to adulthood accompanied with dramatic changes in physical and mental characteristics as well as in social environments^1^. Perhaps because of these overarching changes, adolescence is particularly vulnerable to a range of distress that often leads to psychological and behavioral problems, and, in extreme cases, to psychiatric diseases^2-4^. Sexual maturation and dimorphism are significant biological factors behind the adolescent psychopathology. For instance, boys are more associated with externalizing problems characterized by aggression and rule-breaking than girls, who are often susceptible to internalizing problems such as depression and withdrawal^5-7^. Understanding the risk factors and neural mechanisms underlying these adolescent problems is a critical step towards the identification of modifiable intervention targets to foster health and development of children^8^.

Among other possible targets, the child-parent relationship has been recognized as a crucial environmental factor that impacts on adolescent’s cognition, emotion, and behaviors^9,10^. Therefore, the effect of child-parent relationship on brain development and its relation to adolescent behaviors have been of central importance in adolescent neuroscience and epidemiology^11^. In epidemiological literature, the quality of child-parent relationship has been shown to significantly impact on internalizing and externalizing problems in boys and girls^12^. Furthermore, paternal or maternal relationship with child has been suggested to have differential effects on adolescent behaviors between boys and girls. For instance, the relationship with mother has a stronger influence on boy’s externalizing problems than that with father^3^. On the other hand, the significance of the relationship with father on girl’s perilous sexual behavior has been recently recognized^13^. Together with the well-documented evidence for sexual differences in adolescent brain development^14-17^ and behavioral problems^6,7^, these studies indicate that father and mother’s parenting may differently contribute to specific psychobehavioral problems, which may further interact with child’s sex.

Neuroscientific investigations are expected to reveal the intermediate mechanism underlying the complicated relationship between parenting and child’s psychobehavioral problems^18,19^. Although important evidence for the effect of parenting on the structural brain development in school-age children and adolescents is accumulating ^20-22^, possible differences in paternal and maternal influences, as well as child’s sex differences, remain unexamined. In this regard, a previous resting-state fMRI (rs-fMRI) study of early adolescents^23^ that examined the development of inter-regional functional connection (FC) marks a notable exception. In the study, insensitive parent caregiving, maternal rather than paternal in particular, accelerated the development of FC between the amygdala and medial prefrontal cortex, a key component in the neural circuit for emotion regulation underlying a range of child’s psychobehavioral problems^24,25^. Furthermore, this effect was dependent on the child’s sex, such that the effect was more pronounced in girls than boys. A significant effect of maternal parenting on the development of this particular FC was further shown by another fMRI study that found an accelerated development by early maternal deprivation^26^. However, recent evidence indicates that emotion regulation is subserved by multiple FC networks including the fronto-parietal (FP) and default-mode networks (DMN) ^27,28^, and the FC involving the amygdala may be a part embedded in one such large-scale FC network. Therefore, a whole-brain connectome-based analysis, rather than analysis of specific FCs of interest, may be necessary for the complete understanding of the functional development of neural circuits^29,30^.

Although the association between parenting and the child’s development of FCs has been addressed in several studies, it has been often left unanswered whether and how such FC networks are actually associated with adolescent psychobehavioral problems. Importantly, one seminal longitudinal study addressed this question and showed that maternal aggressive parenting in early adolescence had an effect on the rs-FCs seeding amygdala in mid adolescence, which had a mediation effect on the development of major depressive disorder in late adolescence^31^. However, the critical question of sexual dimorphism in adolescent behavioral problems, as well as possible differential association of paternal- and maternal-child relationships with a child’s behavior, remains unknown. Although functional imaging studies have traditionally adopted correlation methods to show the association between brain measure (e.g. rs-FC) and behavior, the conceptual advantage of a more robust approach based on statistical prediction has been increasingly recognized in neuroimaging literature^32^. However, the possible sex-dependent association between psychobehavioral problems and FC networks related to paternal- and maternal-child relationships has not yet been examined even by correlation analysis.

This study aimed to reveal the mappings between child-parent relationships, child’s FCs, and child’s psychobehavioral problems by considering the differences between paternal and maternal relationships as well as child’s sexual dimorphism in FC and psychobehavioral problems. Here, we have used an epidemiological and neuroimaging dataset of early adolescents derived from the Tokyo TEEN Cohort (TTC) study, which is a population-based birth cohort study in Japan. Our approach had the following two major steps. First, based on possible interaction of paternal or maternal parenting with child’s sex on child’s problems, we performed connectome-based analyses using children’s rs-fMRI data to identify separate functional brain connectomes (FBCs) associated with paternal- and maternal-child relationships, each of which was further divided into the presence or absence of interaction effect with a child’s sex. We then built connectome-based predictive models (CPMs)^33^ for child’s psychobehavioral problems using the identified FBCs to test possible associations between the FBCs and the child’s problems. Together with subsequent control analyses and a causal discovery method, we aimed to reveal the mechanism linking the child-parent relationship, brain, and psychobehavioral problems in early adolescents.

## Results

We analyzed a cohort dataset of 93 early adolescents (41 boys and 52 girls) who were between 10 and 13 years old at the time of data collection (see Table 1 for the demographic information). The main data consisted of the rs-MRI data and questionnaires of the Network Relationship Inventory (NRI) for assessing the child-parent relationship^34,35^ and the Child Behavior Checklist (CBCL) for assessing the psychobehavioral problems in adolescents^36^. See Materials and Methods and Supplementary Information for details.

**Table 1.** Demographic information of children included in this study.

|  | Sex |  | Statistics |  |  |  |
| --- | --- | --- | --- | --- | --- | --- |
|  | Boy (41) | Girl (52) | df | t-value | P-value | Total (93) |
| Age (years) | 12.12 ± 0.80 | 11.84 ± 0.76 | 91 | 1.70 | 0.09 | 11.96 ± 0.79 |
| Estimated IQ<sup>a</sup> | 112.54 ± 17.24 | 107.94 ± 11.86 | 91 | 1.52 | 0.13 | 109.97 ± 14.58 |
| Socioeconomic status | 4.93 ± 0.69 | 4.62 ± 0.84 | 91 | 1.92 | 0.06 | 4.76 ± 0.79 |
| Median FD [mm] | 0.10 ± 0.02 | 0.09 ± 0.03 | 91 | 0.54 | 0.59 | 0.10 ± 0.03 |
| **NRI scores for father** |  |  |  |  |  |  |
| Companionship | 3.60 ± 0.90 | 3.45 ± 1.02 | 91 | 0.02 | 0.98 | 3.51 ± 0.97 |
| Reassurance of worth | 3.94 ± 0.80 | 3.89 ± 1.04 | 91 | -0.37 | 0.71 | 3.92 ± 0.94 |
| Secure base | 4.23 ± 0.75 | 4.03 ± 1.00 | 91 | -0.11 | 0.91 | 4.12 ± 0.90 |
| **NRI scores for mother** |  |  |  |  |  |  |
| Companionship | 3.77 ± 0.81 | 3.86 ± 0.88 | 91 | -0.33 | 0.74 | 3.82 ± 0.85 |
| Reassurance of worth | 4.08 ± 0.74 | 4.08 ± 0.91 | 91 | -0.20 | 0.84 | 4.08 ± 0.83 |
| Secure base | 4.44 ± 0.61 | 4.32 ± 0.86 | 91 | 0.28 | 0.78 | 4.38 ± 0.76 |
| **NRI scores for friend** |  |  |  |  |  |  |
| Companionship | 3.77 ± 0.74 | 3.90 ± 0.68 | 91 | -0.87 | 0.38 | 3.84 ± 0.71 |
| Reassurance of worth | 3.11 ± 0.85 | 3.63 ± 0.76 | 91 | -3.10 | **<0.01** | 3.40 ± 0.84 |
| Secure base | 3.26 ± 0.97 | 3.75 ± 0.79 | 91 | -2.65 | **0.01** | 3.54 ± 0.90 |
| **Psychobehavioral problems<sup>b</sup>** |  |  |  |  |  |  |
| **Internalizing problems** |  |  |  |  |  |  |
| internalizing problem | 51.48 ± 7.37 | 53.82 ± 7.98 | 87 | -1.43 | 0.16 | 52.74 ± 7.75 |
| withdrawn/depressed | 54.35 ± 4.95 | 56.78 ± 6.05 | 87 | -2.05 | **0.04** | 55.66 ± 5.67 |
| somatic complaints | 53.91 ± 5.97 | 52.48 ± 3.70 | 87 | 1.38 | 0.17 | 53.14 ± 4.90 |
| anxiety/depressed | 53.44 ± 5.18 | 54.97 ± 5.38 | 87 | -1.36 | 0.18 | 54.26 ± 5.31 |
| **Externalizing problems** |  |  |  |  |  |  |
| externalizing problem | 50.62 ± 6.52 | 50.58 ± 7.27 | 87 | 0.03 | 0.98 | 50.60 ± 6.90 |
| rule-breaking behavior | 54.78 ± 4.41 | 53.83 ± 5.37 | 84 | 0.89 | 0.38 | 54.26 ± 4.95 |
| aggressive behavior | 52.95 ± 4.24 | 53.22 ± 4.68 | 87 | -0.28 | 0.78 | 53.10 ± 4.46 |
| **Other subscales** |  |  |  |  |  |  |
| social problem | 53.73 ± 4.37 | 54.50 ± 5.26 | 87 | -0.74 | 0.46 | 54.15 ± 4.86 |
| thought problem | 51.56 ± 2.68 | 54.59 ± 7.56 | 87 | -2.44 | **0.02** | 53.20 ± 6.01 |
| attention problem | 53.90 ± 4.93 | 55.60 ± 6.19 | 87 | -1.42 | 0.16 | 54.82 ± 5.68 |
| **Abbreviations:** FD: frame-wise displacement, IQ: intelligence quotient, and NRI: network of relationship inventory. |  |  |  |  |  |  |
| All the scores were described as mean ± standard deviation (SD). |  |  |  |  |  |  |
| <sup>a</sup> Full-scale IQ was estimated using WISC-III. |  |  |  |  |  |  |
| <sup>b</sup> The severity of psychobehavioral problems were measured by the child behavior checklist. The subscale score of rule-breaking behavior was missing for two boys and five girls, respectively. For other subscales, scores were missing for four girls. |  |  |  |  |  |  |

The overview of the analytical procedures is shown in Fig. 1. First, we constructed individual functional connectome using a brain parcellation scheme based on the Brainnetome Atlas^37,38^ from the rs-fMRI in each subject. Second, we identified multiple sets of FCs, each of which was associated with a different aspect of child-parent relationships. Third, we built predictive models for child’s psychobehavioral problems using the identified sets of FCs.

**Fig. 1.**
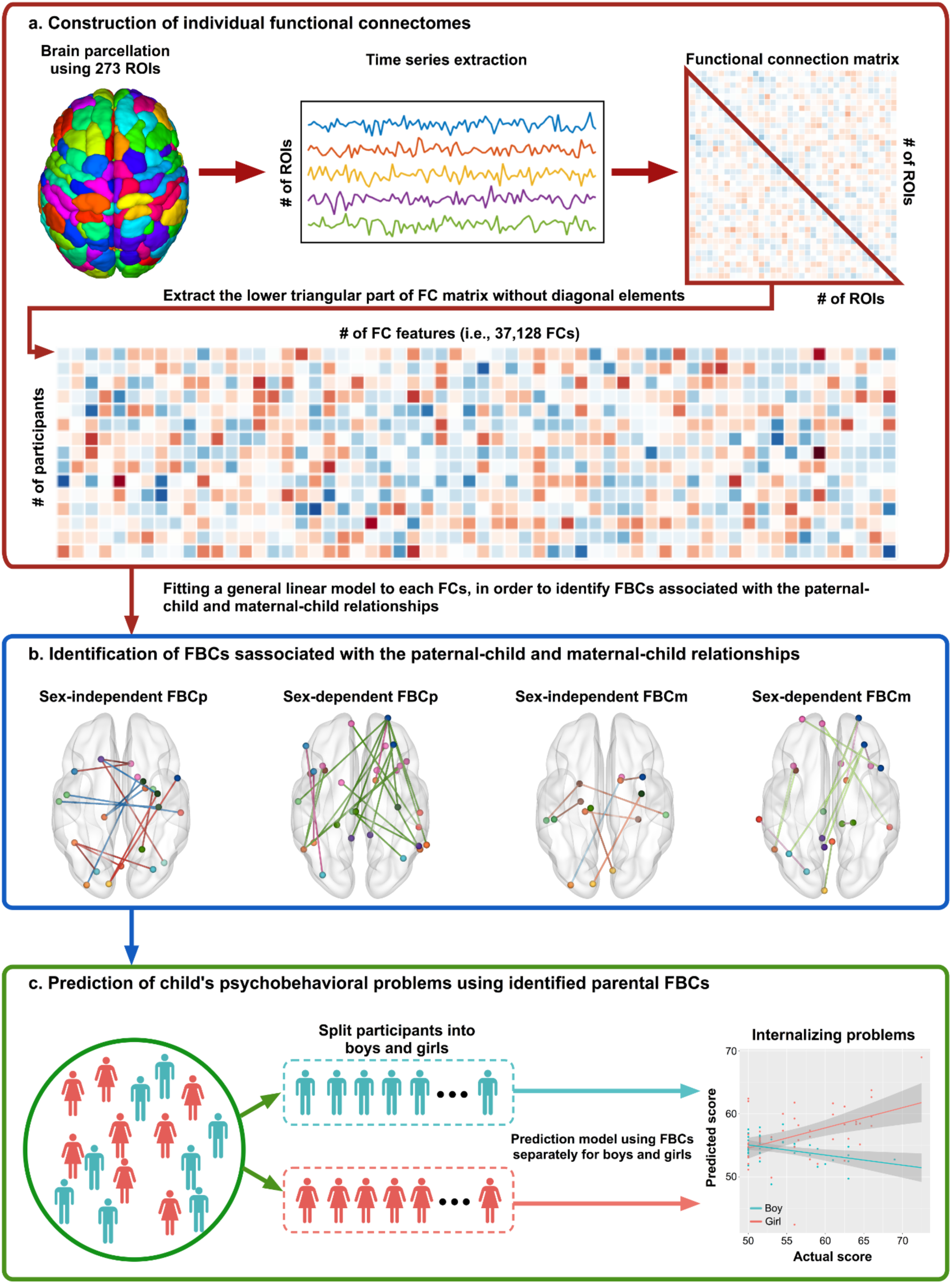
Overview of analytical procedures in this study. Individual functional connectome was constructed using 273 regions of interest (ROIs)^37,38^, each element of which represented a strength of functional connection (FC) between a pair of two ROIs. The lower triangular part of the 273 × 273 FC matrix without diagonal elements was extracted and stored as a row vector of feature matrix. The dimension of the row vector was 37,128 (a). A general linear model was fitted to each of FCs to examine the effects of paternal- and maternal-child relationships separately. Functional brain connectomes (FBCs) associated with the effect of paternal- or maternal-child relationship, which we termed FBCp or FBCm, respectively, were further divided into child’s sex-independent and sex-dependent effects. Statistical threshold was set to *P* < 0.05, false discovery rate (FDR) corrected (b). For prediction of child’s psychobehavioral problems, participants were divided into two groups of boys and girls. Then, prediction models for child’s psychobehavioral problems using these parental FBCs were built by adopting connectome-based prediction (CPM) approach with leave-one-subject-out cross-validation (LOOCV) (c). Prediction accuracy was evaluated by calculating the Pearson correlation coefficient between predicted and actual scores of psychobehavioral problems. Statistical significance was set to *P* < 0.05.

After constructing the individual functional connectomes, each FC was fitted with two general linear models using either variables of the paternal-child relationship or those of the maternal-child relationship. These models allowed us to separately identify sets of FCs that were associated with paternal or maternal relationships. Each of the “paternal” or “maternal” model included explanatory variables for effect of the child-parent relationship and those for *interaction* of child-parent relationship with the child’s sex. The NRI scores for fathers and mothers were used as variables representing the degree of positive relationships with father and with mother in the models, respectively.

### FBCs associated with the paternal-child relationship (FBCp) and maternal-child relationship (FBCm)

Figure 2 and Table S1 show FBCs of the adolescent brain that were significantly associated with the relationship with father or mother, which we termed FBCp and FBCm, respectively. Each of the FBCs was divided into sex-independent (Figs. 2a and 2b) and dependent (Figs. 2c and 2d) FBCs based on the presence or absence of interactions with the child’s sex. Sex-independent FBCp or FBCm further had “positive” and “negative” sub-FBCs with the former increasing its FCs with the NRI score regardless of the child’s sex and the latter vice versa (Fig. 2e). The positive FBCp consisted of nine inter-network FCs, while the negative FBCp contained six FCs (Fig. 2a). On the other hand, positive FBCm contained seven inter-network FCs, while the negative FBCm comprised one FC (Fig. 2b). Comparisons between the FBCp and FBCm revealed only two common FCs: an FC between the right precentral gyrus (PrG) and left lateral occpital (LO) and the other between left LO and right basal ganglia (BG) (see Fig. S1a).

**Fig. 2.**
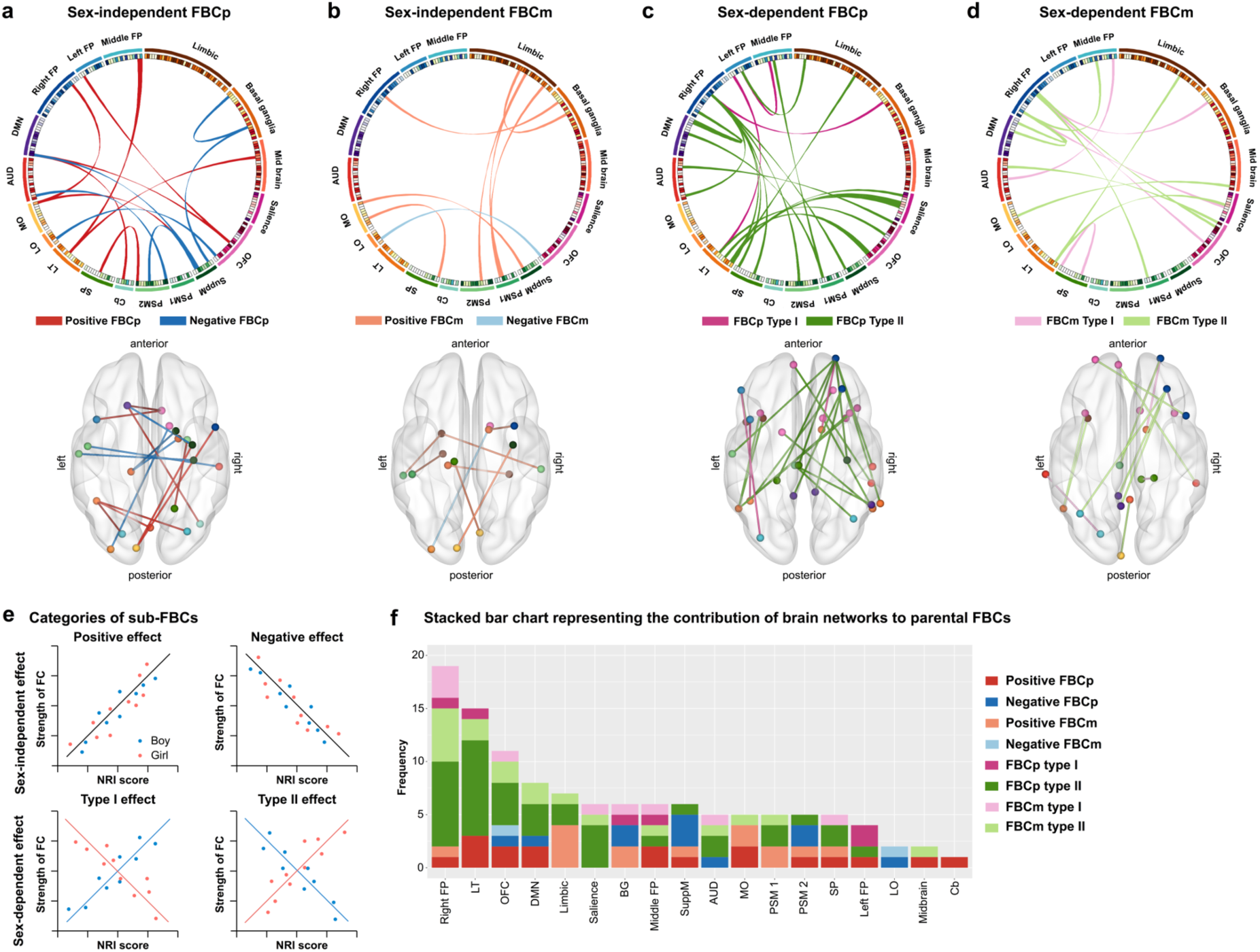
Depictions of functional brain connectomes (FBCs) associated with the paternal- and maternal-child relationships and their characteristics. The connectograms represent FBCs associated with the paternal-(FBCp) and maternal-child (FBCm) relationships along with the corresponding brain maps below each connectogram. FBCs associated with paternal- and maternal-child relationships are divided into two major FBCs exhibiting sex-independent (a, b) and sex-dependent (c, d) effects, each of which are further subdivided into two sub-FBCs exhibiting positive and negative effects. The red and blue ribbons represent the positive and negative effects for sex-independent FBCs, respectively, while the magenta and green ribbons represent the different interaction effects for sex-dependent FBCs. Illustrative examples of these effects are given in (e). The contribution of each brain network defined based on a previous meta-analytic study^39^ to these FBCs are shown in (f).

### Child’s sex-dependent parental FBCs

We identified sets of FCs that showed different directions of associations with child-parent relationship between boys and girls. When the relationships with father or mother showed such a pattern, we termed such FCs as sex-dependent FBCp or FBCm, respectively (Fig. 2e). As shown in Fig. 2 and Table S1, sex-dependent FBCp contained 23 FCs particularly involving the FP, lateral temporal (LT), orbitofrontal (OFC), salience networks, and DMN (Fig. 2c). FBCm consisted of fourteen inter-network connections between the right FP, OFC, salience networks, and DMN (Fig. 2d). The sex-dependent FBCp and FBCm shared five FCs mainly stemming from the right FP to other networks, such as the DMN and BG networks, and from the LT to limbic and salience networks (see Fig. S1b). These observations indicate that, although some commonalities do exist, paternal- and maternal-child relationships are associated with largely different sets of FCs. The presence of abundant FCs in these FBCs indicates that relationships with father and mother interact with child’s sex at the neuronal level.

### Spatial distributions and functional characterization of parental FBCs

In order to characterize the distribution of FCs in the FBCp and FBCm, we classified nodes of FCs in each type of the FBCs. As shown in Fig. 2f, we observed that the right FP network had the largest number of nodes showing any effects of child-parent relationship, most notably interaction effects with the child’s sex. This observation suggests strong sex-dependent influences of the child-parent relationships on the development of this functional network.

In order to further examine functional characteristics of these FBCs, we used the BrainMap (http://brainmap.org)^39^ and examined the spatial overlaps of these FBCs with brain activity patterns associated with brain functional terms registered in the BrainMap. We show a radar chart of each FBC depicting the functional profile as defined in the five most superordinate terms of the BrainMap (Fig. S2 and Table S2; see Table S3 for the results of all mental subdomains). On comparison between FBCp and FBCm, it is notable that functional profiles of sex-dependent FBCp and FBCm are strongly associated with “cognition” and “emotion”. Therefore, paternal and maternal relationships may be particularly related to a child’s cognitive and emotional functions in a sex-dependent manner.

### Prediction of child’s psychobehavioral problems using parental FBCs

Next, we examined whether these parental FBCs were associated with psychobehavioral problems of adolescent child. As a means of testing the association, we constructed CPMs using these FBCs to predict child’s psychobehavioral problems and tested the prediction accuracies of the models. Because previous epidemiological studies showed sex differences in the manifestations of the child’s internalizing and externalizing problems^12^, as well as in the effect of child-parent relationship on psychobehavioral problems^3^, we split the cohort into the two groups based on child’s sex and built separate predictive models for each sex. Each model used eight measures of the FBCs (4 sub-FBCs per FBCp and FBCm. See **Identification of FBCs showing the child’s sex-independent and dependent effects of child-parent relationships** in **Material and Methods**) as predictor variables and one of the subscale scores of the CBCL as a dependent variable. We focused on the CBCL subscales that were related to either internalizing or externalizing problems (see Methods for more details). We used a leave-one-subject-out cross-validation (LOOCV) to evaluate the prediction accuracies of the models, in which correlation between predicted and observed score of a CBCL subscale was calculated. As shown in Fig. 3a, models accurately predicted the psychobehavioral problems in a highly sex-dependent manner: aggressive behavior, one of the externalizing problems, was significantly predicted for boys [*r* = 0.304, *P* = 0.027, *df* = 39] but not for girls. In contrast, subscales related to internalizing problems, such as withdrawn/depressed [*r* = 0.459, *P* < 0.001, *df* = 46], somatic problem [*r* = 0.302, *P* = 0.019, *df* = 46], anxiety/depressed [*r* = 0.431, *P* = 0.001, *df* = 46], and internalizing problem [*r* = 0.534, *P* < 0.001, *df* = 46] were significantly predicted only for girls. For the results of other subscales, see Fig. S3.

**Fig. 3.**
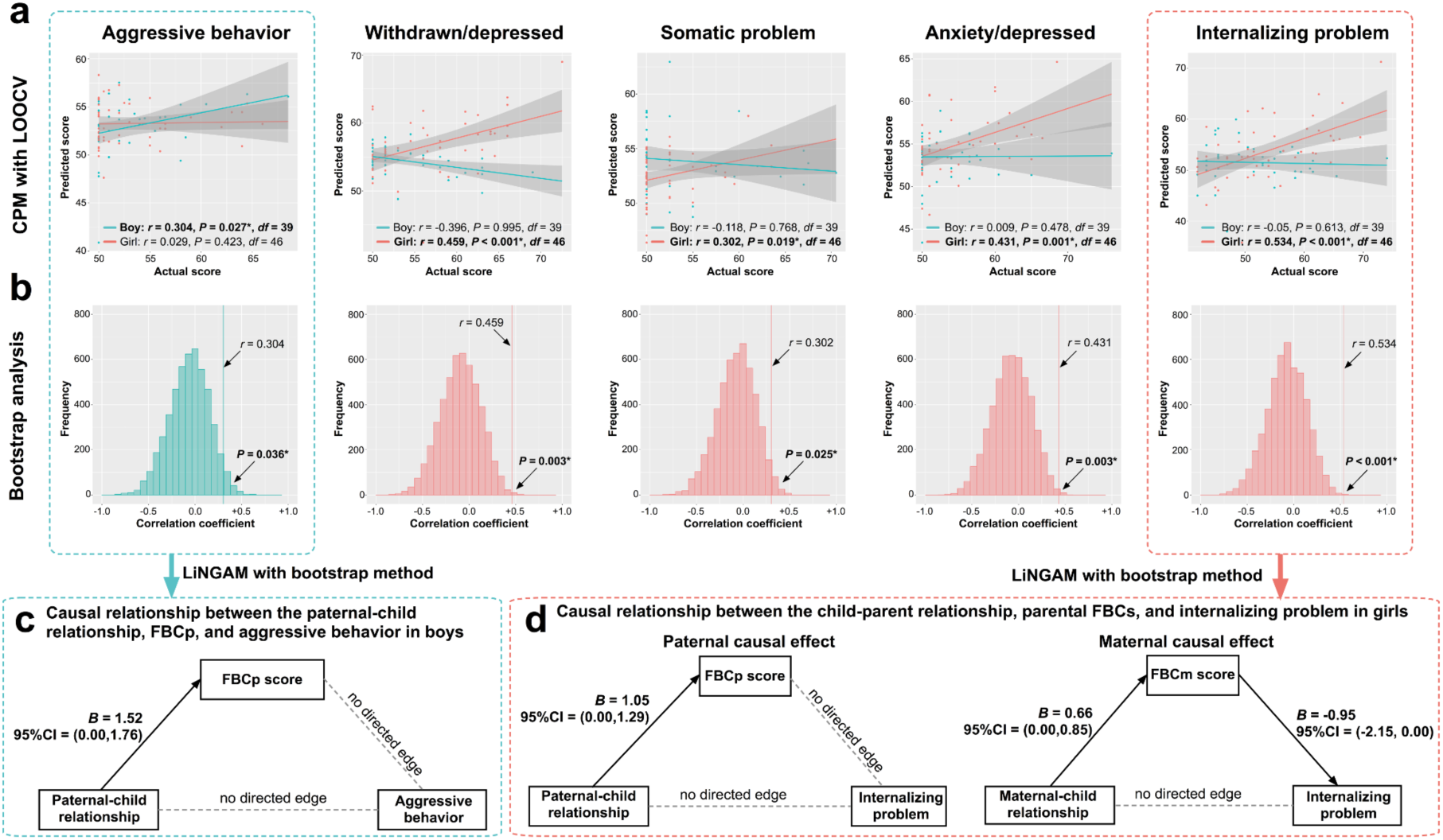
Association of child’s externalizing and internalizing problems with parental functional brain connectomes (FBCs) The scatter plots show association between actual versus predicted scores of child’s externalizing and internalizing problems by prediction models using parental FBCs (a). Prediction models for child’s externalizing and internalizing problems were built using independent variables of parental FBCs. The pink and green dots stand for data points of girl and boy, respectively. Prediction models exhibited statistically significant prediction performance in a highly sex-dependent manner: aggressive behavior for boys [*r* = 0.304, *P* = 0.027, *df* = 39] and subscales related to internalizing problems, such as withdrawn/depressed [*r* = 0.459, *P* < 0.001, *df* = 46], somatic problem [*r* = 0.302, *P* = 0.019, *df* = 46], anxiety/depressed [*r* = 0.431, *P* = 0.001, *df* = 46], and internalizing problem [*r* = 0.534, *P* < 0.001, *df* = 46] for girls. The histograms show null distributions of prediction performance obtained by a bootstrap method with 5,000 iterations (b). LiNGAM^46^ with bootstrap method was applied to discover causal paths between the child-parent relationship, FBCs, and externalizing and internalizing problems in boys and girls. For aggressive behavior in boys, LiNGAM revealed causal path from the paternal-child relationship to FBCp, but other edges were not retained (c). For the internalizing problem in girls, paternal causal effects were only observed in a causal path from the paternal-child relationship to FBCp, while maternal causal effects were observed in causal paths from the maternal-child relationship to FBCm and from FBCm to the internalizing problem (d).

### Bootstrapping analysis to test associations between parental FBCs and psychobehavioral problems

Although the analyses using the LOOCV procedure showed significant prediction accuracy for child’s psychobehavioral problems, one might raise a concern about the possibility of “overfitting” in the prediction models^40^. In order to examine this possibility, we performed a bootstrapping analysis, in which prediction performances were compared with models that were built using randomly selected FCs not included in either type of FBCp or FBCm. Using a bootstrapping analysis with 5,000 iterations ^41^, the Pearson’s correlation values of the prediction model of FBCp and FBCm were compared with the null distribution derived from the bootstrapping procedure. As shown in Fig. 3b, statistical significance of the model using all the parental FBCs was shown in boy’s aggressive behavior (*P* = 0.036) and in girl’s withdrawn/depressed (*P* = 0.003), somatic problem (*P* = 0.025), anxiety/depressed (*P* = 0.003), and internalizing problems (*P* < 0.001). For other subscales, see Fig. S4. These results show that prediction accuracies in the main analyses using the LOOCV procedure were indeed significant. As an additional confirmation, we used permutation tests as well. We confirmed that the results of the permutation tests exactly matched with those of the bootstrap method (see Fig. S5).

### Prediction of psychobehavioral problems using either of FBCp or FBCm

A next important question would be to ask which of the two, FBCp or FBCm, was associated with these internalizing and externalizing problems. We, thus, built prediction models using either of the FBCp or FBCm. Of note, each of the FBCs included both sex-independent and dependent FBCs. As shown in Fig. S6, the models using the FBCp exhibited significant prediction for the subscale of aggressive behavior in boys [*r* = 0.335, *P* = 0.016, *df* = 39] and for the subscale of internalizing problem in girls [*r* = 0.258, *P* = 0.038, *df* = 46]. On the other hand, the models using FBCm significantly predicted girl’s withdrawn/depressed [*r* = 0.449, *P* < 0.001, *df* = 46], anxiety/depressed [*r* = 0.314, *P* = 0.015, *df* = 46], and internalizing problem [*r* = 0.424, *P* = 0.001, *df* = 46] subscales (Fig. S7). These results suggest that FBCp is associated with a part of boy’s externalizing problem, the aggressive behavior, and the internalizing problem in girls, while FBCm is predominantly associated with a range of girl’s internalizing problems.

### Prediction performance of models that do not account for differences in child’s sex or in parental FBCs

Although we built separate prediction models for child’s psychobehavioral problems, these psychobehavioral problems might be more accurately and parsimoniously predicted without assuming separate models for boys and girls. To examine this possibility, we built a single prediction model for each CBCL subscale combining the data of boys and girls. As shown in Fig. S8, we observed statistically significant correlations in several psychobehavioral problems: aggressive behavior [*r* = 0.241, *P* = 0.012, *df* = 87], internalizing problem [*r* = 0.186, *P* = 0.040, *df* = 87], and externalizing problem [*r* = 0.205, *P* = 0.027, *df* = 87]. However, overall predictive performance was degraded including withdrawn/depressed [*r* = 0.086, *P* = 0.210, *df* = 87], somatic complaints [*r* = 0.052, *P* = 0.314, *df* = 87], and anxiety/depressed [*r* = 0.166, *P* = 0.060, *df* = 87], all of which were accurately predicted in the separate model for girls. We noted that the prediction performance degraded despite clear advantage of the combined analysis regarding the data size in calculating correlation. These observations are consistent with the view of sex-dependent effects of paternal and maternal FBCs on child’s psychobehavioral problems.

Another possible complexity of our prediction model might be that the effects of the paternal- and maternal-child relationship on FCs were examined separately. If, however, role of the paternal- and maternal-child relationship was not distinct but interchangeable, then FBCs associated with the combined measure of both parents could also predict child’s psychobehavioral problems. To examine this possibility, we defined the child-parent relationship as the sum of the NRI scores of both parents. We repeated the analyses to identify sex-independent and dependent FBCs associated with the combined measure of child-parent relationships, based on which we built prediction models for the CBCL subscales (see Supplementary Information for further details). However, the models failed to predict any CBCL subscale scores (Fig. S9). These results indicate that psychobehavioral problems in child are better predicted by modeling differential effects of FBCp and FBCm than by assuming comparable or interchangeable effects.

### Prediction of psychobehavioral problems by models using FBCs associated with peer relationships, family SES, and psychobehavioral problem itself

In addition to the child-parent relationship, the relationship with peers is thought to increase importance in adolescent’s life^42^. It is, thus, interesting to ask whether FBCs associated with the peer relationship are related to psychobehavioral problems in children. Therefore, we repeated the series of analyses to identify FBCs associated with the peer relationship and constructed prediction models for psychobehavioral problems using the identified FBCs (see Supplementary Information for further details). The degree of positive relationship between child and his/her peers was assessed by the NRI. Prediction models using FBCs associated with the child-peer relationship could not predict any CBCL subscales (all *P* > 0.120; Fig. S10). These results suggest that, at least for early adolescence, the child-parent relationship, rather than that the peer relationship, is an important factor associated with child’s behavioral outcomes. We also confirmed that FBCs associated with family SES, another important factor associated with adolescent psychobehavioral problems^43,44^, could not predict any CBCL subscales (Fig. S11; see Supplementary Information for further details).

In the main analyses adopting the CPM approach, we successfully predicted key psychobehavioral problems of boys and girls from the FBCp and FBCm. However, one may be concerned that a significant prediction might be achieved simply because the predicted behavioral variable (i.e., a CBCL subscale score) might be correlated with another behavioral variable (i.e., NRI score) associated with FBCs. In order to examine this possibility, we next compared performance of prediction in the main analysis with that of the CPM model to predict child’s psychobehavioral problem using FBCs associated with that psychobehavioral problem. Here, the predicted behavioral variable and the variable associated with FBC are exactly the same, which would produce high prediction accuracy, if the correlation between predicted variable and FBC-associated variable is critical. We built prediction models for each psychobehavioral problem using a standard CPM approach as described in previous studies^33,45^. As shown in Fig. S12, these analyses showed statistically significant correlation between predicted and actual values only in boy’s attention problem [*r* = 0.420, *P* = 0.003, *df* = 39] but not in any of the subscales related to internalizing and externalizing problems. These results suggested that the FBCp and FBCm measures are actually significant predictors for the behavioral symptoms in adolescents.

### Causal discovery among child-parent relationship, FBC, and psychobehavioral problems

Our main and control analyses demonstrated that the paternal- and maternal-child relationships were associated with sets of FBCs, that were further associated with specific psychobehavioral problems in adolescents. Although we have examined the chain of the triad associations among child-parent relationships, brain, and child’s problems in a hypothesis-driven manner, we further questioned whether the causal paths between the three components could be inferred in a data-driven manner. For this purpose, we used the Linear Non-Gaussian Acyclic causal Model (LiNGAM)^46,47^. The methodological details can be found elsewhere^46^. Based on a previous study^46^, a bootstrap method with 10,000 iterations was applied to compute the 95% confidence interval (CI) of connection strengths. We retained the direct edges only if the 95% CI did not include null value (i.e., zero) (see Methods for details).

Because the CBCL “aggressive behavior” scale of boy was the only scale belonging to the externalizing problem that the parental FBCs significantly predicted, we focused on this scale as an instance of the externalizing problem. For the internalizing problem, however, we focused on the CBCL “internalizing problem” subscale of girl because it is a superordinate metric over other related subscales, such as withdrawn/depressed, somatic problem, and anxiety/depressed, that the parental FBCs significantly predicted.

#### Boy’s Aggressive Behavior

Because the FBCp, but not FBCm, predicted the severity of boy’s aggressive behavior, we incorporated father-related measures (i.e., composite score of FBCp and the NRI score for father) into the analysis. These inputs satisfied the assumption of non-Gaussianity (see Table S4). Of note, *P*-values were derived from the one-sample Kolmogorov-Smirnov (KS) tests and the interval (*x, y*) stands for the open interval between *x* and *y*. As shown in Fig. 3c, LiNGAM with bootstrap method revealed the direct causal effect from the paternal-child relationship to the FBCp [*B* = 1.52, 95% CI = (0.00, 1.76)], while other direct edges did not survive.

#### Girl’s Internalizing Problem

The CPM analyses indicated that both FBCp and FBCm significantly predicted the severity of internalizing problems in girls, although the latter showed better prediction than the former. Therefore, we examined the causal graphs of both father-related measures (composite score of FBCp and NRI score for father) and mother-related measures (composite score of FBCm and NRI score for mother). We confirmed that these variables satisfied the assumption of non-Gaussianity (see Table S4). Although the maternal composite score and the severity of internalizing problem included null value within the interval, the one-sample KS tests confirmed that these measures were far from the normal distribution. LiNGAM with bootstrap method revealed the direct causal effect from the girl-father relationship to the brain [*B* = 1.05, 95% CI = (0.00, 1.29)] in the model for paternal causal effect. On the other hand, in the model for maternal causal effect, we observed the direct causal effect from the girl-mother relationship to the FBCm [*B* = 0.66, 95% CI = (0.00, 0.85)], and from the FBCm to the severity of internalizing problems [*B* = −0.95, 95% CI = (−2.15, 0.00)] (Fig. 3d).

## Discussion

The current study analyzed a cohort dataset of early adolescents to identify the pattern of the rs-FCs of the child’s brain that are associated with the child-parent relationship and further to test whether and how multiple FBCs associated with distinct types of child-parent relationships contribute to child’s psychobehavioral problems. First, our general linear models separately identified effect of paternal and maternal relationships on FCs, which we termed FBCp and FBCm, respectively. We, then, built CPMs using these FBCs to predict psychobehavioral problems of boys and girls separately. We observed a sex-dependent pattern in prediction performance of the models, such that a part of externalizing problems was significantly predicted only for boys whereas the internalizing problems were predicted only for girls. Further analyses revealed that FBCp was associated with a part of boy’s externalizing problem and girl’s internalizing problem, whereas FBCm was associated with a range of girl’s internalizing problems. Series of control analyses confirmed that the present model best predicted child’s problems among other models that did not account for differences in child’s sex or in paternal and maternal relationships. Furthermore, similar connectome-based predictive models using effect of the peer relationship or the family SES on FBCs did not predict child’s problems either. Therefore, our hypothesis-driven approach demonstrated the sex-dependent associations between the child-parent relationships, FBCs, and child’s psychobehavioral problems in early adolescence. Parts of the casual relationships among these components were specified by causal discovery, which indicated the causal paths from the quality of girl-mother relationship to FBCm, and further to girl’s internalizing problems.

Effect of child-parent relationship on adolescent’s brain development has been examined in previous structural and functional neuroimaging studies^18,20,21^. However, many of them focused on the effect of maternal parenting, and only one study has examined the possible differences between paternal or maternal relationship on child’s brain development or interaction of child-parent relationship with child’s sex^25^. To the best of our knowledge, the present study is the first to separately associate paternal or maternal relationships with child’s FCs in a sex-dependent manner using the functional connectome-based analysis. We observed that FCs included in FBCp and FBCm were unevenly distributed among major FC networks, with FCs involved in the right FP network showing particularly strong associations (Fig. 2). The spatial distribution of these FCs was highly different from those FCs associated with the peer relationship or family SES (Fig. 4). This observation may indicate distinct effect of child-parent relationship on child’s brain among other environmental factors influencing on adolescents’ behavior. Comparison between FBCp and FBCm revealed marked differences between the two FBCs, although several common FCs did exist. Together with the observation that a notable number of FCs in each of the two FBCs showed child’s sex-dependent associations with child-parent relationship, our findings indicate that the paternal-child and maternal-child relationships are differently associated with FCs in the child’s brain, which are further different between boys and girls.

**Fig. 4.**
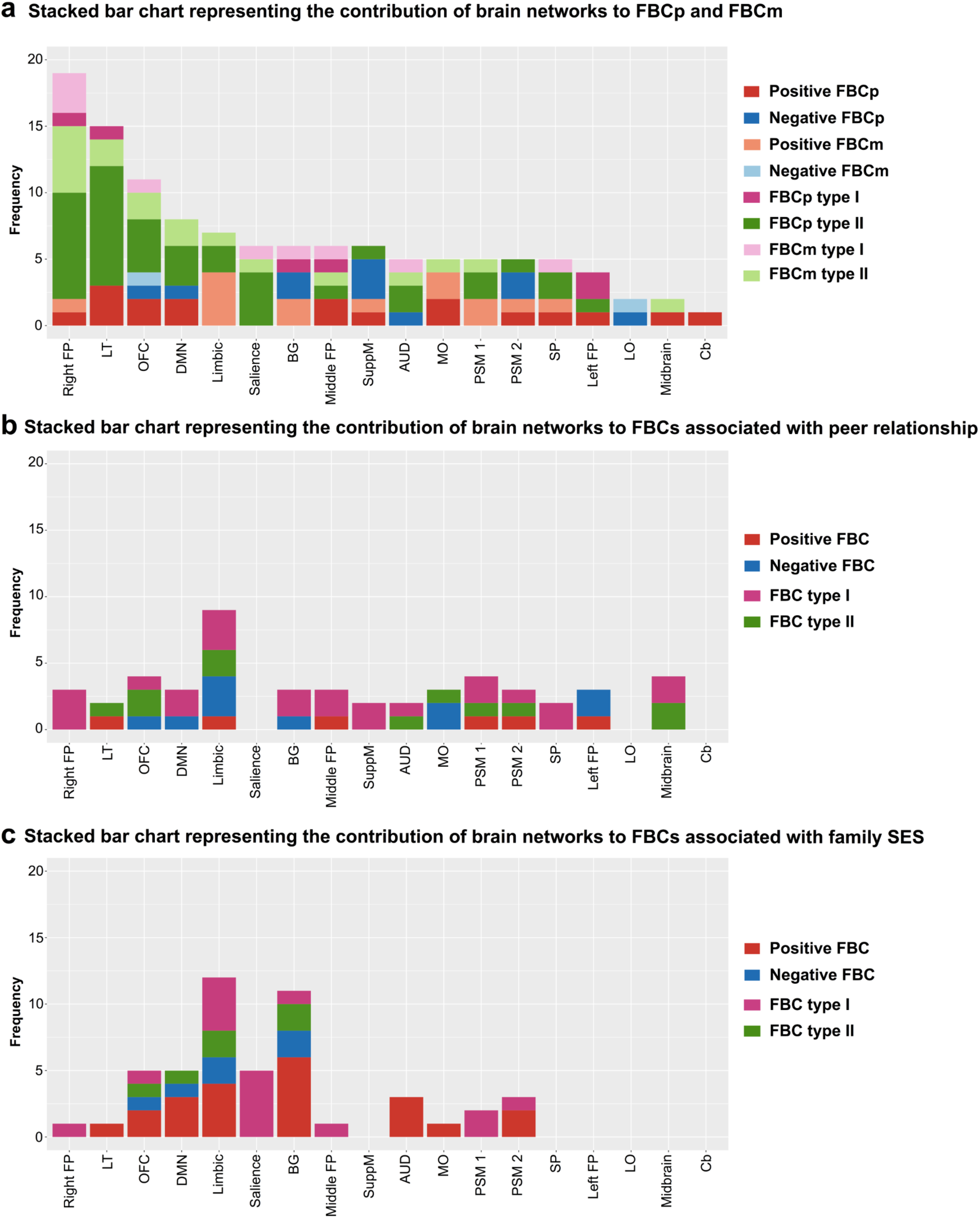
Stacked bar charts representing the contribution of brain networks to FBCs associated with either of the child-parent relationships, the peer relationship, or family SES. The top stacked bar chart represents the contribution of brain networks defined by a previous meta-analytic study to FBCs associated with both of the child-parent and maternal-child relationships. The child-parent relationship affects strongly on the right fronto-parietal (FP) network. The middle-stacked bar chart represents the contribution of brain networks to FBCs associated with the peer relationship. The peer relationship affects mainly on the limbic network. The bottom stacked bar chart shows the contribution of brain networks to FBCs associated with family SES. Family SES has an effect mainly on the limbic and basal ganglia (BG) networks.

The clear sex difference in association of FCs with child-parent relationship deserves further discussion. Our observation of sex-dependent association between child-parent relationship and FCs is consistent with the previous finding of child sex-specific effect of parenting on brain development^21,23^. In particular, we found that the large portion of FCs with sex-dependent effect of child-parent relationship were presented in the right FP network. This finding is consistent with a previous structural MRI study reporting that the effect of maternal negative parenting on the cortical thinning in the right FP regions is different between adolescent boys and for girls^21^. Interestingly, deficit of executive functions subserved in the right FP network have been associated with a gamut of behavioral problems^50^. Such considerations highlight the development of executive functions as a crucial factor that interacts with emotion and social functions for adolescent’s well-being and adaptive behaviors^51^. It is possible that sex-dependent effect of child-parent relationship on the neural systems including the right FP network is particularly critical for the development of cognition and emotion functions that are required for regulation of adaptive and flexible behavior^51^, failure of which may lead to a range of sex-dependent behavioral problems in adolescence.

Previous adolescent functional imaging studies examining parenting effects on child’s FCs were motivated by the notion that parenting-related FCs would contribute to child’s functional outcomes^25,26^. However, the triplet relation of parenting, FCs, functional outcomes has not been demonstrated except for one longitudinal study showing the impact of maternal aggressive parenting in early adolescence on amygdala-seeded FCs in mid-adolescence, which had a mediation effect on major depression disorder in late adolescence^31^. To the best of our knowledge, our study is the first demonstration of the tripartite link by building predictive models for sex-dependent child’s psychobehavioral problems using measures of FBCs associated with paternal-child and maternal-child relationships. Suprisingly, the models showed significant prediction in a highly sex-dependent manner, such that the model accurately predicted a part of externalizing problems in boys and the range of internalizing problems in girls. We further showed different association between FBCp and FBCm, with the former being associated with boy’s externalizing problems and girl’s internalizing problems and the latter being associated with girl’s internalizing problems. Although our cross-sectional study design did not allow the assumption regarding the causal relationship among child-parent relationship, FCs, and psychobehavioral problems based on temporal order, the data-driven analysis of LiNGAM indicated the casual paths from the relationship with mother to FBCm, and from FBCm to internalizing problems in girls. Taken together, these observations indicate different roles of paternal and maternal parenting on an adolescent’s brain development and further suggest sex-dependent mechanisms that link parenting, brain, and psychobehavioral outcomes.

It is notable that child’s psychobehavioral problems were best predicted when models treated measures of FBCp and FBCm independently and when boy’s and girl’s psychobehavioral problems were modeled separately. This observation again supports the importance of the interaction between child-parent relationship and child’s sex in understanding of the child’s functional outcomes. We also demonstrated that FCs associated with the relationship with peers^12,52^ or family SES^43,44^ failed to predict any of the psychobehavioral problems, although effect of these factors on adolescent’s functions and behaviors have been recognized. Given that the relationship with peers may increase importance towards the adulthood, it is possible that these factors might play more important roles later in adolescence and that the relationship with parents is more important in influencing child’s brain development and psychobehavioral outcomes^8^ at least in early adolescence.

In conclusion, the present study revealed a differential association of paternal and maternal relationships with adolescent’s FBCs, which was further different between boys and girls. CPMs incorporating the FBCp significantly predicted boy’s externalizing and girl’s internalizing problems, whereas the model incorporating the FBCm predicted girl’s internalizing problems. Our findings, thus, provide sex-dependent neuronal substrates underlying the epidemiological observations of complex relation between paternal and maternal parenting and psychobehavioral problems in adolescents.

## Materials and Methods

### Cohort dataset

Data were provided by a longitudinal cohort study (TTC; http://ttcp.umin.jp/) for adolescence in the Tokyo area. Main data used in this study consisted of the three components: questionnaires for (i) child-parent relationship and (ii) psychobehavioral problems of adolescent children and (iii) MRI data of adolescent children. In addition, basic information of the child and his/her family, including intelligence quotient (IQ) and socioeconomic status (SES), was collected through a semi-structured interview by trained interviewers. Further details of data collection procedures in TTC are described elsewhere^53,54^. This study was approved by the ethics committees of the University of Tokyo, the Tokyo Metropolitan Institute of Medical Science, and the Tokyo Metropolitan University.

For the assessment of the child-parent relationship, we used the NRI^34,35^, in which an adolescent child rated the degree of positive relationship with his/her father and mother separately. For the assessment of psychobehavioral problems in children, we used the CBCL^36^, answered by the child’s primary caregiver. This study focused on the CBCL items that were relevant to the internalizing and externalizing problems. In addition to these measures, the child’s full-scale IQ and family’s SES were estimated using the Wechsler Intelligence Scale for Children third edition and a brief rating scale of SES^55^, respectively, at the TTC survey for a 10-year-old adolescent. These measures were included as a nuisance covariate in the general linear model when identifying the FBCs associated with the paternal- or maternal-child relationships (see **Identification of FBCs showing the child’s sex-independent and dependent effects of child-parent relationships**).

In this study, we selected 93 children (41 boys and 52 girls) from the TTC dataset. The details of participant selection procedures are described in Supplementary Information. Table 1 shows the demographic information of the participants included in the present analysis.

### MRI acquisition

The MRI data were collected using 3.0-T Philips Achieva scanner (Philips Medical Systems, Best, The Netherlands) with an 8-channel receive head coil. Functional images were acquired using echo planar imaging sequence (repetition time [TR]: 2.5 s, echo time [TE]: 30 ms, flip angle: 80°, 3.2 mm thickness with a 0.8-mm gap, field of view [FoV]: 212 mm, matrix size: 64 × 64, 40 slices, 250 volumes). During the resting scan, children were asked to fixate their gaze on a cross-hair displayed on the screen. An anatomical image was also acquired using the MPRAGE sequence (TR: 7.0 ms, TE: 3.2 ms, flip angle: 9°, 1 mm slice thickness, FoV: 256 × 240 mm, matrix size: 256 × 240, 200 sagittal slices, voxel size: 1 × 1 × 1 mm).

### Rs-fMRI processing

All the functional images were preprocessed using Statistical Parametric Mapping 12 (SPM12; Wellcome Department of Cognitive Neurology, London, UK) with a subsidiary use of in-house scripts in the following steps: 1) slice timing and head motion correction, 2) co-registration of functional images to anatomical image, 3) normalization and resampling to 2 mm isotropic voxel using the DARTEL^56^, and 4) spatial smoothing using a Gaussian kernel (6-mm FWHM). We excluded participants if their head motion was larger than 3 mm or 3° from the first volume. For each participant, the median frame-wise displacement (FD) was calculated from six head motion parameters to quantify the amount of head motion during scans^57^. The mean and standard deviation of median FD are shown in Table 1. We further excluded participants whose median FD was greater than 0.15 mm, as previously recommended^33^.

To remove the effects of head motions during scans, we used ICA-AROMA^58,59^ and further performed nuisance signal regression^60^. Nuisance signals consist of signals extracted from white matter, cerebrospinal fluid, and gray matter binary masks^61^. A band-pass filter (0.008 – 0.1 Hz) was then applied. Individual functional connectome was constructed using 273 regions of interest (ROIs) provided by the Brainnetome atlas^37^ combined with the cerebellum atlas^38^. For each participant, a 273 × 273 correlation matrix was calculated using the Pearson correlation coefficient among all possible pairs of ROIs. Each element of this matrix was then converted to *z*-score using Fisher’s *r*-to-*z* transformation.

### Identification of FBCs showing the child’s sex-independent and dependent effects of child-parent relationships

To identify FCs that were associated with the child-parent relationship, the following general linear model was fitted to each FC:

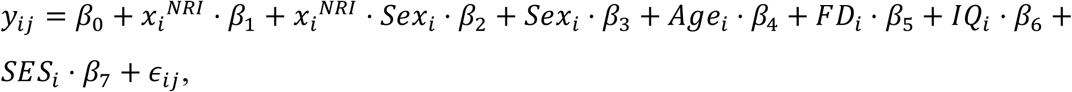

where *y*_*i.j*_ stands for the *j*-th FC of the *i*-th participant and *x*_*i*_^*NRI*^ stands for either father’s or mother’s NRI score for the *i*-th participant. Age (*Age*_*i*_), frame-wise displacement of head (*FD*_*i*_), child’s IQ (*IQ*_*i*_), and SES (*SES*_*i*_) were included as additional nuisance covariates. Here, we would highlight that two different models were fitted to each FC in order to independently assess the effects of relationships with father and mother. Furthermore, the effect of the relationship with father or mother on child’s FCs were divided into two variables of child’s sex-independent term (β_1_) and the sex-dependent (interaction) term (β_2_). Therefore, this analysis identified four major sets of FCs (i.e., FBCs): 1) the FBCs showing the sex-independent effects of paternal relationship, 2) the ones showing the sex-independent effects of maternal relationship, 3) the ones showing the sex-dependent effects of paternal relationship (interaction of father’s NRI score and child’s sex), and 4) the ones showing the sex-dependent effects of maternal relationship. Each major FBC was further subdivided into two sub-FBCs depending on the directions of association between FC and NRI score (i.e., positive and negative effects), resulting in eight FBCs in total (see **Prediction of child’s psychobehavioral problems from FBCs associated with child-parent relationship**). Statistical significance was set to *P* < 0.05 corrected for false discovery rate (FDR) implemented in the Network-Based Statistics toolbox^62^.

### Prediction of child’s psychobehavioral problems from FBCs associated with child-parent relationship

We used a method of CPM^33,45^ with LOOCV to examine whether the FBCs identified by the previous steps were predictive of psychobehavioral problems of children. We split the participants into two groups based on sex and built a separate prediction model for each group. In order to apply the CPM with LOOCV, we first calculated connectivity strengths for the eight types of FBCs (see **Identification of FBCs showing the child’s sex-independent and dependent effects of child-parent relationships**), as to each participant: (1a) summation of FCs positively associated with the paternal-child relationship (“sex-independent positive FBCp”), (1b) summation of FCs negative associated with the paternal-child relationship (“sex-independent negative FBCp”), (2a) summation of FCs positively associated with the maternal-child relationship (“sex-independent positive FBCm”), (2b) summation of FCs negatively associated with the maternal-child relationship (“sex-independent negative FBCm”), (3a) summation of FCs exhibiting an interaction effect between the paternal-child relationship and sex such that the effects of the paternal-child relationship are larger for boys than girls (“sex-dependent FBCp type I”), (3b) summation of FCs exhibiting an interaction effect between the paternal-child relationship and sex such that the effects of the paternal-child relationship are larger for girls than boys (“sex-dependent FBCp type II”), (4a) summation of FCs exhibiting an interaction effect between the maternal-child relationship and sex such that the effects of the maternal-child relationship are larger for boys than girls (“sex-dependent FBCm type I”), and (4b) summation of FCs exhibiting an interaction effect between the maternal-child relationship and sex such that the effects of the maternal-child relationship are larger for girls than boys (“sex-dependent FBCm type II”). A prediction model was built using these eight summarized variables.

For the CPM with LOOCV, an iterative two-step analytical procedure was performed as follows: 1) building of a predictive model using *N*-1 training data, and 2) testing on the left-out participant. Here, multiple linear regression was used to build a predictive model where predictor variables were the eight summarized network measures described above and the dependent variable was a CBCL subscale score of interest. The constructed CPM was then used to predict a score of the subscale for the left-out participant. The predictive power of this model was assessed by the Pearson correlation coefficient between predicted and actual scores across boys or girls separately. Due to the explorative nature of this analyses, statistical threshold was set to *P* < 0.05, uncorrected.

### Causal discovery among child-parent relationships, FBCs, and psychobehavioral problems

A series of analyses confirmed the existence of triad associations among the child-parent relationship, parental FBCs, and child’s psychobehavioral problems. In order to examine the causal paths between them, we applied a statistical algorithm for causal discovery. Standard mediation analysis makes explicit assumptions about causal directions beforehand, which may not be appropriate for this study that adopts the cross-sectional design. Here, we used a recently developed data-driven method called the Linear Non-Gaussian Acyclic causal Model (LiNGAM) for causal discovery. LiNGAM is a non-Gaussian variant of structural equation modeling^47^ and several variants of LiNGAM have been applied in various fields^48,49^. In this study, we used a direct estimation algorithm named DirectLiNGAM^46^. Briefly, this algorithm first evaluates pairwise independence between a variable and each of residuals to find an exogenous variable. Next, it removes the effect of the exogenous variable from the other variables using least squares regression. This algorithm repeats these steps until all the causal ordering are determined. The methodological details can be found elsewhere^46^. The MATLAB code for DirectLiNGAM used in this study is available on the website (https://sites.google.com/site/sshimizu06/Dlingamcode). Based on a previous study^46^, a bootstrap method with 10,000 iterations was applied to compute the 95% confidence interval (CI) of the causal path. We retained the direct edges only if the 95% CI did not include null value (i.e., zero).

As in the CPM analyses, separate models were constructed for boys and girls. We chose the CBCL subscale of “aggressive behavior” as boy’s psychobehavioral problem in the model because this was the only subscale that was significantly predicted from parental FBCs. On the other hand, we focused on the CBCL subscale “internalizing problems” as girl’s psychobehavioral problem in the model because it is a superordinate metric over “withdrawn/depressed”, “somatic problem”, and “anxiety/depressed”, that the parental FBCs significantly predicted.

Although each of the FBCp and FBCm has multiple FBCs (e.g. sex-independent and dependent FBCs with positive and negative effects or with type I and type II interaction), inclusions of all these FBCs as multiple independent variables in the model would decrease the interpretability of the model and statistical powers in estimation of the causal model. Therefore, we introduced the following composite score for the FBCp and FBCm respectively:

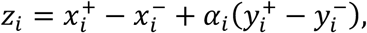

where the two variables, *x*_*i*_ and *y*_*i*_, represent sex-independent FBC and sex-dependent FBC of *i*-th participant, respectively. The superscript, + and −, represent positive and negative effects for *x* (sex-independent FBCs) or type I and type II interaction for *y* (sex-dependent FBCs). *α*_*i*_ stands for the sex of *i*-th participant. If the participant is boy, then *α*_*i*_ = +1. When the participant is girl, then *α*_*i*_ = −1. Hence, the resultant composite score, *z*_*i*_, represents the change of FCs with the goodness of paternal- or maternal-child relationship.

LiNGAM relies on the assumption of non-Gaussianity to discover acyclic causal graph. Thus, we examined whether each input variable hold the properties of non-Guassianity by calculating skewness, *γ*, and kurtosis, *κ*. A bootstrap method with 10,000 iterations was applied to compute the 95% CI. Based on previous studies ^63^, the one-sample Kolmogorov-Smirnov (KS) test was performed to examine the normality of each variable. Statistical threshold was set to *P* < 0.05. All the procedures were performed using *R* software version 3.4.2. As shown in Table S4, we confirmed that all the variables held the properties of non-Gaussianity.

## Acknowledgments

The current was supported by JSPS KAKENHI Grant numbers 16H06395 (K.K.), 16H06396 (R.H.), 16H06398 (A. N.), 16H06399 (K.K.), and 16K21720 (K.K.), Advanced Bioimaging Support Grant number 16H06280 (K.K.); Japan Agency for Medical Research and Development (AMED), grant numbers JP18dm0307001 (K.K.) and JP18dm0307004 (K.K.). The current work was also supported by UTokyo Center for Integrative Science of Human Behavior (CiSHuB), and the International Research Center for Neurointelligence (WPI-IRCN) at the University of Tokyo Institutes for Advanced Study (UTIAS).

