## Supplementary Information for "Functional connectome linking child-parent relationships with psychological problems in adolescence"

**This PDF file includes:**

Supplementary text
Figs. S1 to S16
Tables S1 to S6
References for Supplementary Information

### Supplementary Methods and Results

#### Cohort dataset

All the data used in this study were provided by a longitudinal cohort study (TTC; http://ttcp.umin.jp/) for adolescence in the Tokyo area. Although the TTC is an ongoing study that is currently expanding its dataset, the present study used the subset of the data for early adolescence (10 – 13 years old) that were collected between 2011 and 2017 (First wave: 2012-2015, Second wave: 2014-2017). The details for data collection procedures in TTC are described elsewhere[1,2].

For the assessment of the child-parent relationship, we used the Network Relationship Inventory (NRI)[3,4], in which an adolescent child rated the degree of positive relationships with his/her father and mother separately. As to each parent, three subscales of "companionship," "reassurance of worth," and "seek secure base" were rated and the summation of the three subscales was used as a measure of the paternal-child and maternal-child relationship. In the TTC dataset for early adolescence, the NRI was sampled twice (one at the time of MRI scan [boys: 12.01 ± 0.80 years old, girls: 11.76 ± 0.77 years old, $t$-value = 1.57, $P$ = 0.12, $df$ = 91] and the other at a large-scale survey for 12-year-old adolescents [boys: 12.18 ± 0.30 years old, girls: 12.26 ± 0.30 years old, $t$-value = -1.29, $P$ = 0.20, $df$ = 91]). In order to ensure the reliability and stability of child's assessment, the NRI scores were averaged over the two time-points for analysis.

For the assessment of psychobehavioral problems in children, we used the Child Behavior Checklist (CBCL)[5], answered by the child's primary caregiver. This study focused on the CBCL items that were relevant to the internalizing and externalizing problems. The CBCL subscales related to internalizing problems included: "withdrawal/depressed," "somatic complaints," "anxiety/depressed," and "internalizing problem." On the other hand, the subscales related to externalizing problems included "rule-breaking behavior," "aggressive behavior," and "externalizing problem." Similar to the NRI, the CBCL was also sampled twice (one at a large-scale survey for 10-year-old adolescents [boys: 10.31 ± 0.27 years old, girls: 10.32 ± 0.24 years old, $t$-value = -0.07, $P$ = 0.95, $df$ = 91] and the other at a large-scale survey for the 12-year-old adolescents [boys: 12.18 ± 0.30 years old, girls: 12.26

± 0.30 years old, $t$-value = -1.29, $P$ = 0.20, $df$ = 91]). As was done for the NRI scores, we averaged the scores over the two time-points for the analysis.

**Sex-differences in demographic information**

We performed statistical analyses on the demographic information, including the child-parent relationship, psychobehavioral problems, IQ, and family SES, to examine whether there were sex-related differences in demographic information. There were no significant sex-dependent differences in the paternal-child relationship [boy: 11.80 ± 2.75, girl: 11.90 ± 2.77, $t$-value = -0.17, $P$ = 0.86, $df$ = 91], the maternal-child relationship [boy: 12.57 ± 2.25, girl: 12.65 ± 2.60, $t$-value = -0.17, $P$ = 0.87, $df$ = 91], the estimated IQ [boy: 112.54 ± 17.24, girl: 107.94 ± 11.86, $t$-value = 1.52, $P$ = 0.13, $df$ = 91] or in family's SES [boy: 4.93 ± 0.69, girl: 4.62 ± 0.84, $t$-value = 1.92, $P$ = 0.06, $df$ = 91]. In the psychobehavioral problems, there was a statistically significant sex-related difference for "withdrawn/depressed" [boy: 54.35 ± 4.95, girl: 56.78 ± 6.05, $t$-value = -2.05, $P$ = 0.04, $df$ = 87]. For other variables, see Table 1.

**Participant selection procedure**

In the TTC cohort for early adolescence, 148 children were identified as those with all the necessary data items of the MRI, NRI, and the basic demographic data (FIQ and SES). Because the first step in our analysis was to identify the effect of parental relationships on FC, availability of all data items listed above was the necessary condition for inclusion in the study. From these, we selected the sample for current analysis in the following manner: 1) we first excluded participants when they scored "0: I do not have this person" to any NRI items about the relationship with father or mother; 2) we then excluded participants if their anatomical MRI exhibited either brain anomalies or low signal-to-noise ratio due to head motion; and 3) we further excluded participants if their rs-fMRI data exhibited significant head motions during scan (see **Rs-fMRI processing** for exclusion criteria) or their rs-fMRI data were acquired using different scanning parameters. This excluded 55 children from the dataset and the remaining data from 93 children (41 boys and 52 girls; age [mean ± standard deviation]: 11.96 ± 0.79) were analyzed in this study. From this sample, 2 boys and 5 girls were later excluded during the connectome-based predictive model (CPM) analyses because of their missing CBCL data.

**Functional profiles of FBCp and FBCm**

For the functional characterization of the FBCp and FBCm, we examined the spatial relationships of these FBCs with previously established meta-analytic brain maps provided by BrainMap (http://brainmap.org)[6]. BrainMap offers a taxonomy to classify brain activity patterns into one of 61 mental subdomains (e.g., execution, memory, reward, and sexuality), each of which has a representative meta-analytic brain map. In BrainMap, these subdomains are organized into five superordinate categories: action, cognition, emotion, perception, and interoception. In order to obtain functional profiles of the identified FBCs at the most superordinate level, spatial overlaps between meta-analytic maps and each of the FBCs were calculated and then were averaged within each of the five superordinate categories. As shown in Fig. S2 and Table S2, both paternal and maternal FBCs showed strong associations with "Cognition" and "Emotion" in the sex-dependent effects (type II). Another common characteristic may include relatively strong associations with "Perception" in the sex-independent positive effects. As might be expected, no FBCs exhibited strong association with "Interoception". For the results of subdomains, see Table S3. These results show that both sex-dependent FBCp and FBCm are associated with cognitive and emotional functions of a child.

**Predictions of social, thought, and attention problems using parental FBCs**

In the main analyses of CPM using parental FBCs, we focused on prediction of CBCL subscale scores related to externalizing and internalizing problems. Here we show results of the predictive models for the other CBCL subscales, social problem, thought problem, and attention problem using the exact same methods as the CPMs for externalizing and internalizing problems. The results are summarized in Fig. S3. The models using the FBCp and FBCm exhibited significant prediction performance for girl's thought problem [$r = 0.404$, $P = 0.002$, $df = 46$].

We further constructed prediction models using either FBCp or FBCm to examine which of the two parental FBCs contributed significantly towards prediction. The prediction models with FBCp did not [$r = 0.142$, $P > 0.05$, $df = 46$] (see Fig. S6), whereas the models with FBCm exhibited statistically significant prediction [$r = 0.312$, $P = 0.016$, $df = 46$] (see Fig. S7). These results suggest that the FBCm is associated with girl's thought problem, in addition to the range of girl's internalizing problems.

In order to confirm the association between FBCm and girl's internalizing problem, we performed the same bootstrap method described for the externalizing and internalizing problems. As shown in Fig. S4, the $r$ value for girl's thought problem reached a significant level ($P = 0.003$). For other CBCL subscales, see Fig. S4.

**Permutation test for association between the child's psychobehavioral problems and parental FBCs**

In addition to the bootstrapping analysis described in the main text, we also used a permutation test to confirm whether parental FBCs could predict the psychobehavioral problems in children. Similar to the bootstrap method (see **Bootstrapping analysis to test associations between parental FBCs and psychobehavioral problems** in the Results section of the main manuscript), at each iteration, we shuffled the order of dependent variable (i.e., one of child's psychobehavioral problems) to break the true association between FBCs and each CBCL subscale. Then, we built prediction models for each CBCL with LOOCV procedure. We further evaluated the prediction performance by calculating the Pearson's correlation coefficient between the actual and predicted scores. Repeating these procedures generated a null distribution of prediction performance and the statistical significance of actual performance was assessed using this null distribution. Statistical threshold was set to $P < 0.05$. As shown in Fig. S5, permutation tests completely replicated the results of bootstrap analyses (Fig. S4). These results provide a further support for the associations between parental FBCs and child's psychobehavioral problems.

**Identification of FBCs associated with the combined measure of the child-parent relationship and prediction of child's psychobehavioral problems**

In this control analysis, we examined whether FBCs associated with the combined measure of both parents may also predict child's psychobehavioral problems. As has been done in the main analysis, we fitted the following general linear model to each FC:

$$y_{ij} = \beta_0 + {x_i}^{NRI} \cdot \beta_1 + {x_i}^{NRI} \cdot Sex_i \cdot \beta_2 + Sex_i \cdot \beta_3 + Age_i \cdot \beta_4 + FD_i \cdot \beta_5 + IQ_i \cdot \beta_6 + SES_i \cdot \beta_7 + \epsilon_{ij}.$$

Here, we used the sum of the NRI scores of both parents as the combined measure of the degree of positive relationship with parents. As in the main analyses, the effect of the child-parent relationship on FCs was divided into two variables of child's sex-independent term ($\beta_1$) and the sex-dependent (interaction) term ($\beta_2$). Therefore, this analysis identified two major types of FBCs showing the sex-independent or dependent effect of the parental relationship. Each major FBC was further divided into two sub-FBCs depending on directions of association, resulting in the four sub-FBCs in total. Statistical significance was set to $P < 0.05$ corrected for FDR using the Network-Based Statistics toolbox[7].

Figure S13 shows FBCs associated with the combined measure of child-parent relationships. Similar to the main analysis, the number of FCs involved in sex-dependent FBCs was larger than that involved in the sex-independent FBCs, suggesting the existence of strong sex-dependent effect of the child-parent relationships on FCs. For a descriptive purpose, we show the overlap of FCs identified in this analysis with those included in FBCp and FBCm identified in the main analysis (Fig. S14). The FBCs in this analysis shared 17 FCs (positive: 3 FCs, negative: 2 FCs, type I: 2 FCs, and type II: 10 FCs) with FBCp. On the other hand, the FBCs shared 13 FCs (positive: 3 FCs, negative 1 FC, type I: 4 FCs, and type II: 5 FCs) with FBCm.

We built the prediction models using the identified FBCs using the same method as in the main analysis using CPM with LOOCV (Fig. S9). The models did not predict any of the psychobehavioral problems either in boys (all $P > 0.17$) or in girls (all $P > 0.23$). These results indicate that psychobehavioral problems in child were better predicted by modeling different effects of the relationships with father and mother than by assuming comparable and additive effect of parents.

**Identification of FBCs associated with the child-peer relationship and prediction of child's psychobehavioral problems.**

The NRI also contains questions regarding the degree of positive relationship with peers [3]. As has been done for identification of FBCp and FBCm, we fitted a general linear model to each FC and assessed the sex-independent and sex-dependent effects of the peer relationship while sex, age, median FD, child's IQ, and SES were included as nuisance covariates. As in the previous analyses, this analysis identified the two major types of FBCs exhibiting sex-independent or sex-dependent effect of the peer relationship on FCs. Each type of FBC was subdivided into the two sub-FBCs depending on the direction of association, resulting in the four sub-FBCs in total. We set the statistical significance to $P < 0.05$ corrected for FDR using the Network-Based Statistics toolbox[7].

Figure S15 and Table S5 show FBCs associated with the peer relationship. The sex-independent FBCs consisted of eight inter-network connections stemming from limbic and left FP networks to other networks, such as medial occipital (MO), orbitofrontal (OFC), and basal ganglia (BG) networks. On the other hand, the sex-dependent FBCs consisted of network connections stemming from midbrain and fronto-parietal (FP) networks (right FP and middle FP) to other networks, such as limbic and BG networks. We then examined whether FBCs associated with the peer relationships shared FCs with paternal and maternal FBCs. Interestingly, no FCs were shared between parental FBCs and peer-related FBCs. These results suggest that the effect of the peer relationship on adolescent FCs are largely different from those of the paternal- or maternal-child relationships.

We built the prediction models using the identified FBCs in the same fashion as we did in the main analysis using CPM with LOOCV. Results are shown in Fig. S10. The models did not significantly predict any of the psychobehavioral problems either in boys (all $P >$ 0.64) or in girls (all $P > 0.12$).

**Identification of FBCs associated with the family SES and prediction of child's psychobehavioral problems**

In this control analysis, we examined whether FBCs associated with the family SES might predict child's psychobehavioral problems, motivated by evidence that the SES affects the brain development and psychobehavioral functions during adolescence[8,9]. We fitted a general linear model to each FCs to identify FBCs associated with the effect of SES. It should be noted that we included the NRI scores for both parents as additional nuisance covariates to distill FBCs associated with SES. As in the previous analyses, this analysis identified two types of FBCs of exhibiting sex-independent or sex-dependent effect of SES. Each type of FBC was subdivided into two sub-FBCs based on the direction of association, resulting in the four FBCs in total.

Figure S16 and Table S6 show FBCs associated with the family SES. Sex-independent FBCs consisted of 11 connections dominantly between DMN, BG, and limbic networks. On the other hand, sex-dependent FBCs associated with SES contained 17 connections mainly between or within limbic and salience networks. No FCs were shared with parental FBCs.

We built the prediction models for child's psychobehavioral problems using the identified FBCs in the same fashion as we did in the main analysis by means of CPM with LOOCV. Results are shown in Fig. S12. The models did not significantly predict any of the psychobehavioral problems (all $P > 0.288$).

**Prediction of psychobehavioral problems using FBCs associated with psychobehavioral problems themselves**

In this control analysis, we examined whether FBCs associated with psychobehavioral problems could predict child's psychobehavioral problems. This analysis was motivated by a possibility that significant prediction might be achieved in the CPM approach simply when the predicted behavioral variable (i.e., a CBCL subscale score) was correlated with another behavioral variable (i.e., NRI score) associated with FBC. To examine this possibility, we built predictions models for each psychobehavioral problem using the standard CPM approach as described in previous studies[10-12].

Although a pre-defined threshold (i.e., hyperparameter) might be needed for selecting FCs associated with psychobehavioral problem (CBCL subscale) of interest, we used a leave-one-subject-out nested cross-validation to avoid the bias from hyperparameter tuning[13]. Briefly, we first divided a given dataset into training and test datasets for "outer" loop. A grid search automatically determined the optimal threshold for selecting FCs associated with psychobehavioral problem of interest only using training data during the “inner” loop. We next identified FBCs using the optimal threshold. We calculated a summarized measure separately for FBCs exhibiting positive and negative effects. Then, a regression model was built using the two summarized measures obtained from the training data. Finally, the obtained prediction model was applied to the test dataset. It should be noted that, similar to a previous study[11], FCs associated with nuisance covariates, such as age, median FD, and child’s IQ were excluded during feature selection process.

Results of the prediction models are shown in Fig. S12. Statistically significant prediction was achieved only for attention problem in boys [$r$ = 0.420, $P$ = 0.003, $df$ = 39]. These results suggest that the CPM does not significantly predict behavioral outcomes simply when variables for behavioral outcomes and variables that are associated with FBCs are correlated.

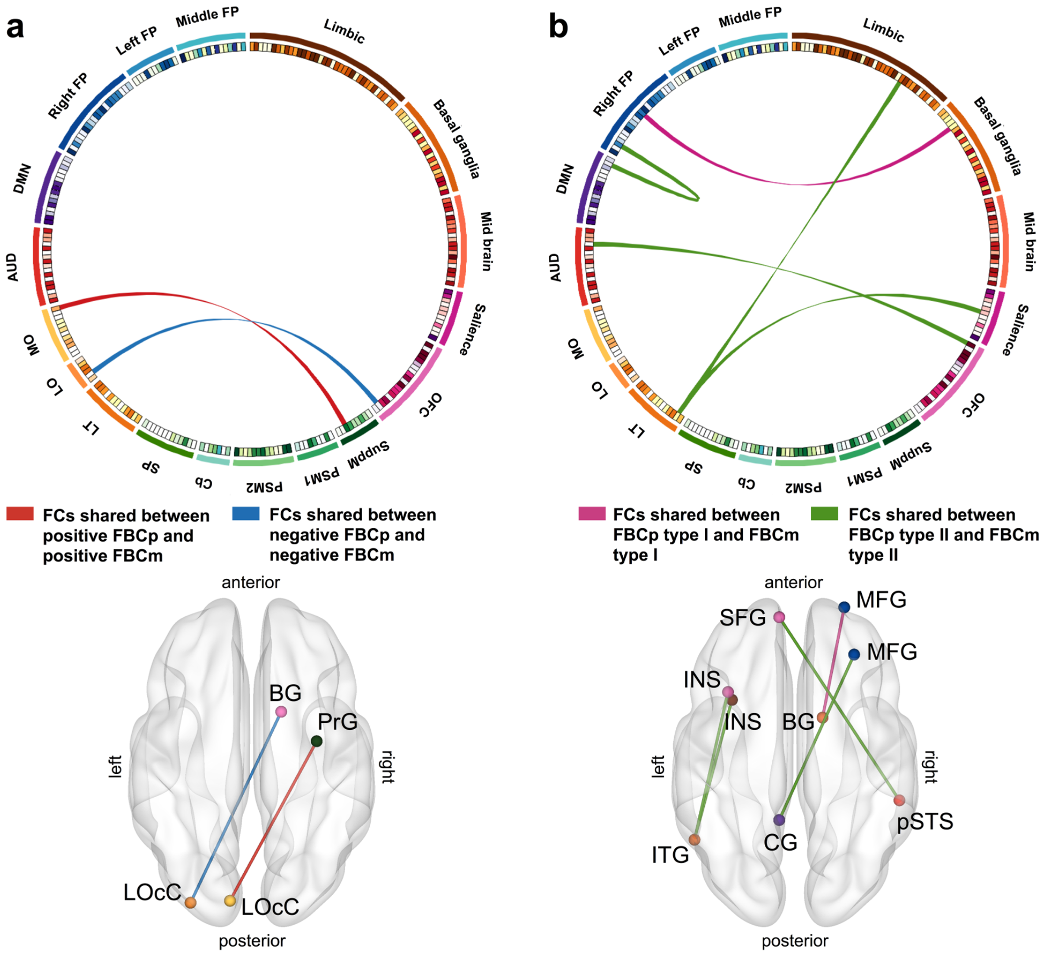


**Fig. S1. Depiction of FCs shared between FBCp and FBCm.**

The left connectogram shows FCs shared between FBCp and FBCm (a). The red and blue ribbons represent an FC shared between positive FBCp and FBCm and the one shared between negative FBCp and FBCm, respectively. The right connectogram shows FCs shared between sex-dependent FBCp and FBCm (b). The pink and green ribbons stand for an FC shared between FBCp type I and FBCm type I and shared FCs between FBCp type II and FBCm type II, respectively.

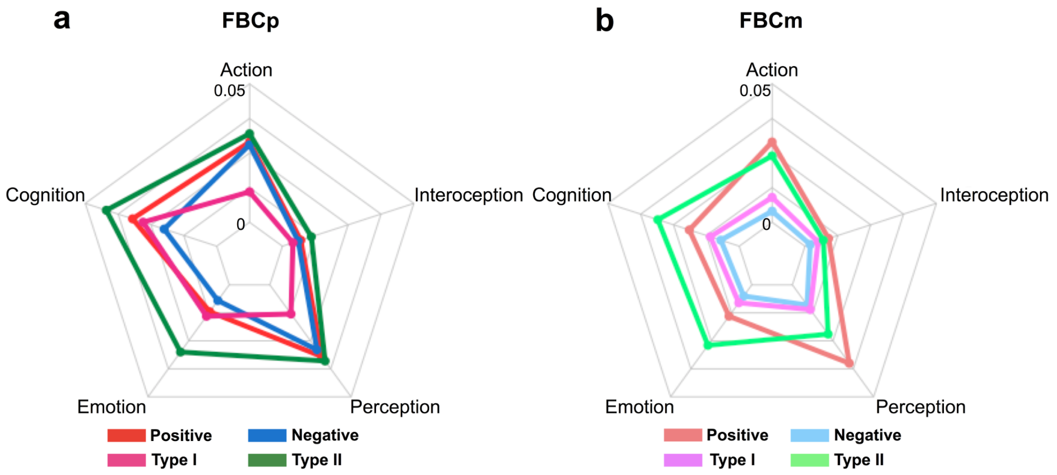


**Fig. S2. Radar charts representing functional profiles of FBCp and FBCm using spatial overlap analyses with meta-analytic maps**

Spatial overlaps between each FBC and meta-analytic map for one of the mental subdomains provided by BrainMap[6] were calculated and then averaged within each superordinate domains: action, cognition, emotion, interoception, and perception. The left radar chart shows the functional profile of FBCp, while the right one shows those of FBCm.

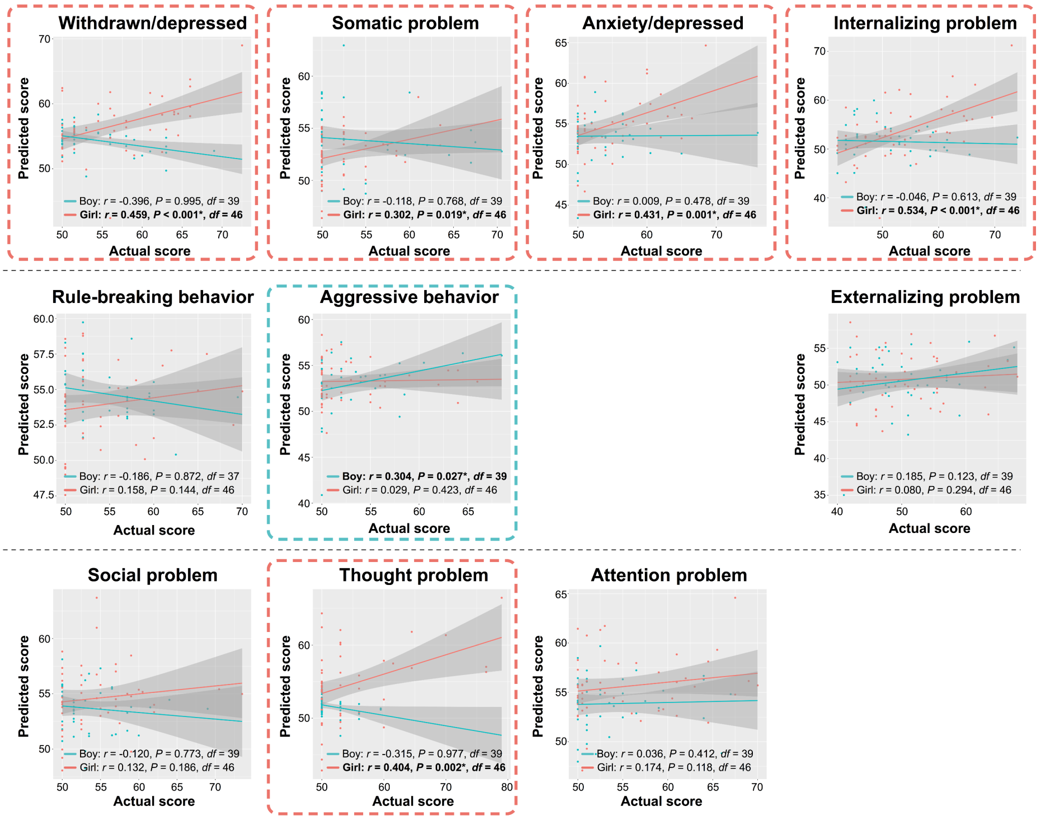


**Fig. S3. Scatter plots of actual versus predicted scores using prediction models with both FBCp and FBCm**

Prediction models for child's psychobehavioral problems were built using independent variables of parental FBCs. The top, middle, and bottom panels show prediction performances for the CBCL subscales of the internalizing problems, the externalizing problems, and other CBCL subscales (social problem, thought problem, and attention problem), respectively. The pink and green dots represent data points of girl and boy, respectively. Statistical threshold was set to $P < 0.05$. Prediction models exhibited significant prediction accuracies for aggressive behavior [$r = 0.304$, $P = 0.027$, $df = 39$] in boy and for withdrawn/depressed [$r = 0.459$, $P < 0.001$, $df = 46$], somatic problem [$r = 0.302$, $P = 0.019$, $df = 46$], anxiety/depressed [$r = 0.431$, $P = 0.001$, $df = 46$], internalizing problem [$r = 0.534$, $P < 0.001$, $df = 46$], and thought problem [$r = 0.404$, $P = 0.002$, $df = 46$] in girl.

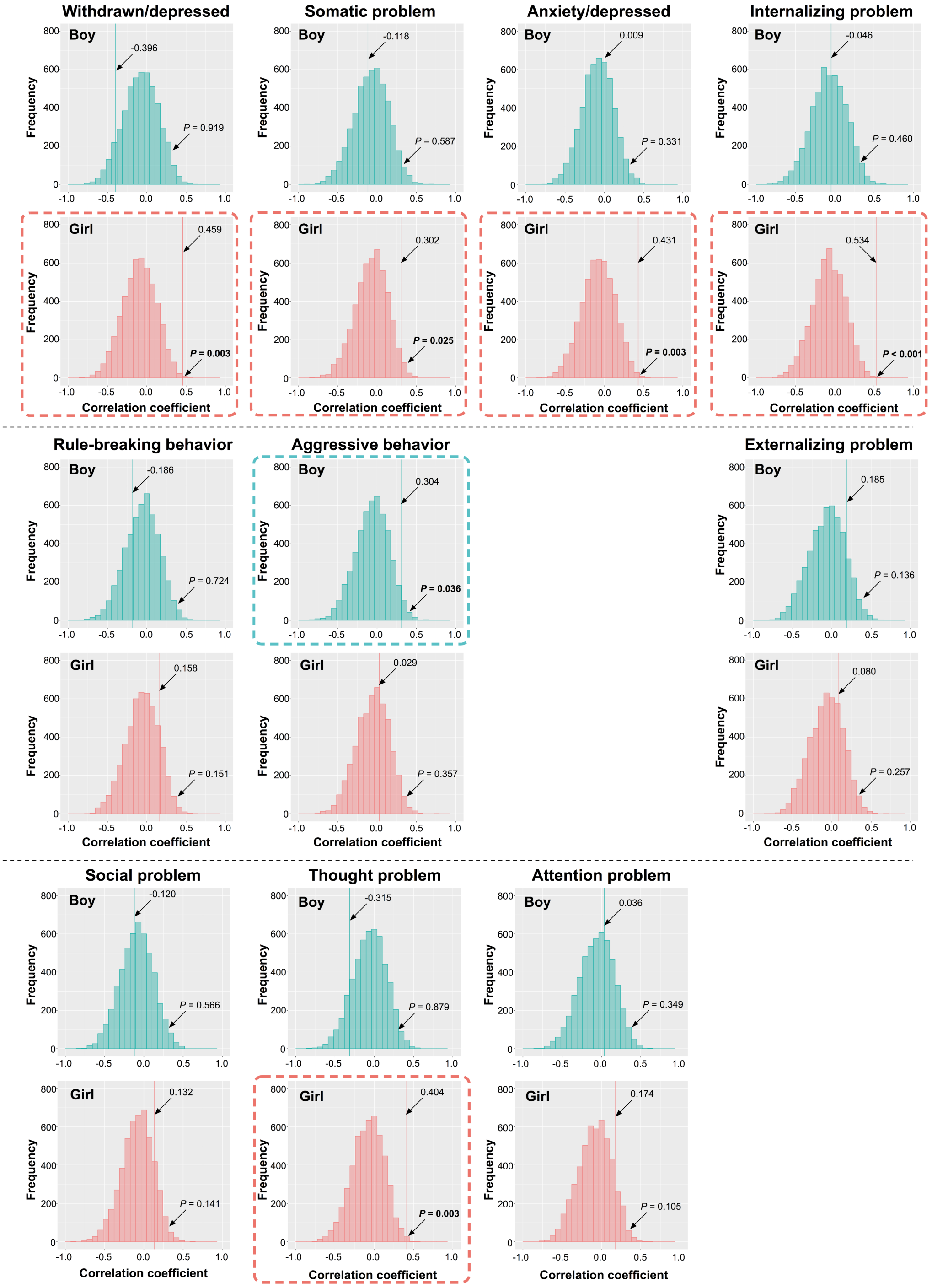
Withdrawn/depressed
Somatic problem
Anxiety/depressed
Internalizing problem
Boy
Girl
-0.396
P = 0.919
-0.118
P = 0.587
0.009
P = 0.331
-0.046
P = 0.460
0.459
P = 0.003
0.302
P = 0.025
0.431
P = 0.003
0.534
P < 0.001
Rule-breaking behavior
Aggressive behavior
Externalizing problem
-0.186
P = 0.724
0.304
P = 0.036
0.185
P = 0.136
0.158
P = 0.151
0.029
P = 0.357
0.080
P = 0.257
Social problem
Thought problem
Attention problem
-0.120
P = 0.566
-0.315
P = 0.879
0.036
P = 0.349
0.132
P = 0.141
0.404
P = 0.003
0.174
P = 0.105
Frequency
Correlation coefficient

**Fig. S4. Histograms of null distribution of prediction performance obtained using a bootstrap method**

Bootstrap method with 5,000 iterations generated the null distribution of prediction performance using randomly selected FCs from the pool of FCs that were not included in either of FBCp or FBCm. The statistical significance of actual performance was assessed using this null distribution. In aggressive behavior, statistical significance was shown only for boy's data ($P = 0.036$). On the other hand, statistical significance was shown for girl's data in withdrawn/depressed ($P = 0.003$), somatic problem ($P = 0.025$), anxiety/depressed ($P = 0.003$), internalizing problem ($P < 0.001$), and thought problem ($P = 0.003$). The results were consistent with Fig. S3.

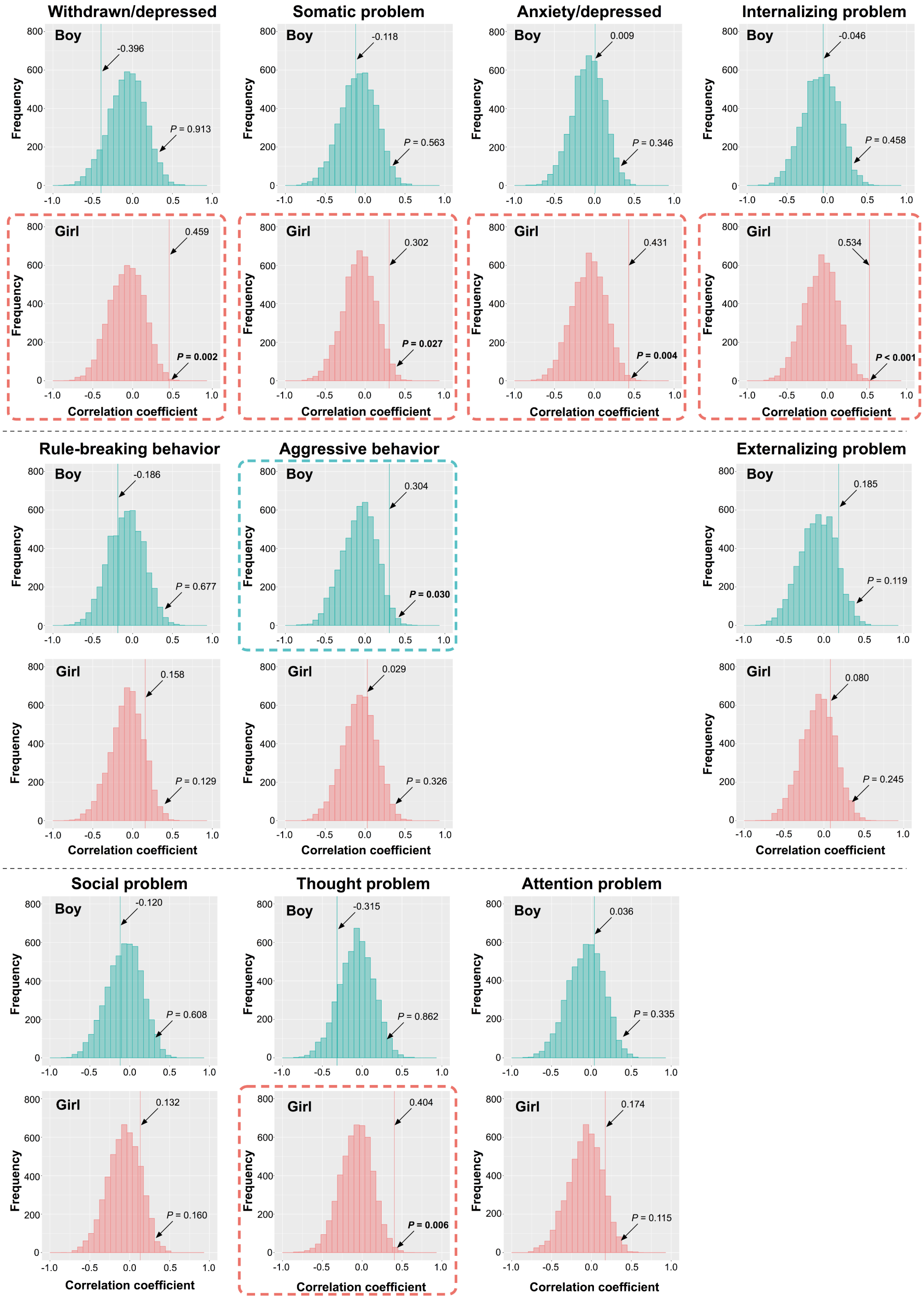
Withdrawn/depressed
Boy
-0.396
P = 0.913
Girl
0.459
P = 0.002
Somatic problem
Boy
-0.118
P = 0.563
Girl
0.302
P = 0.027
Anxiety/depressed
Boy
0.009
P = 0.346
Girl
0.431
P = 0.004
Internalizing problem
Boy
-0.046
P = 0.458
Girl
0.534
P < 0.001
Rule-breaking behavior
Boy
-0.186
P = 0.677
Girl
0.158
P = 0.129
Aggressive behavior
Boy
0.304
P = 0.030
Girl
0.029
P = 0.326
Externalizing problem
Boy
0.185
P = 0.119
Girl
0.080
P = 0.245
Social problem
Boy
-0.120
P = 0.608
Girl
0.132
P = 0.160
Thought problem
Boy
-0.315
P = 0.862
Girl
0.404
P = 0.006
Attention problem
Boy
0.036
P = 0.335
Girl
0.174
P = 0.115
Frequency
Correlation coefficient

**Fig. S5. Histograms of null distribution of prediction performance obtained using permutation test**

Permutation test with 5,000 iterations generated the null distribution of prediction performance using a randomly shuffled data. The statistical significance of actual performance was assessed using this null distribution. In aggressive behavior, statistical significance was shown only for boy's data ($P$ = 0.030). On the other hand, statistical significance was shown for girl's data in withdrawn/depressed ($P$ = 0.002), somatic problem ($P$ = 0.027), anxiety/depressed ($P$ = 0.004), internalizing problem ($P$ < 0.001), and thought problem ($P$ = 0.006). The results matched with Fig.S3 and Fig. S4.

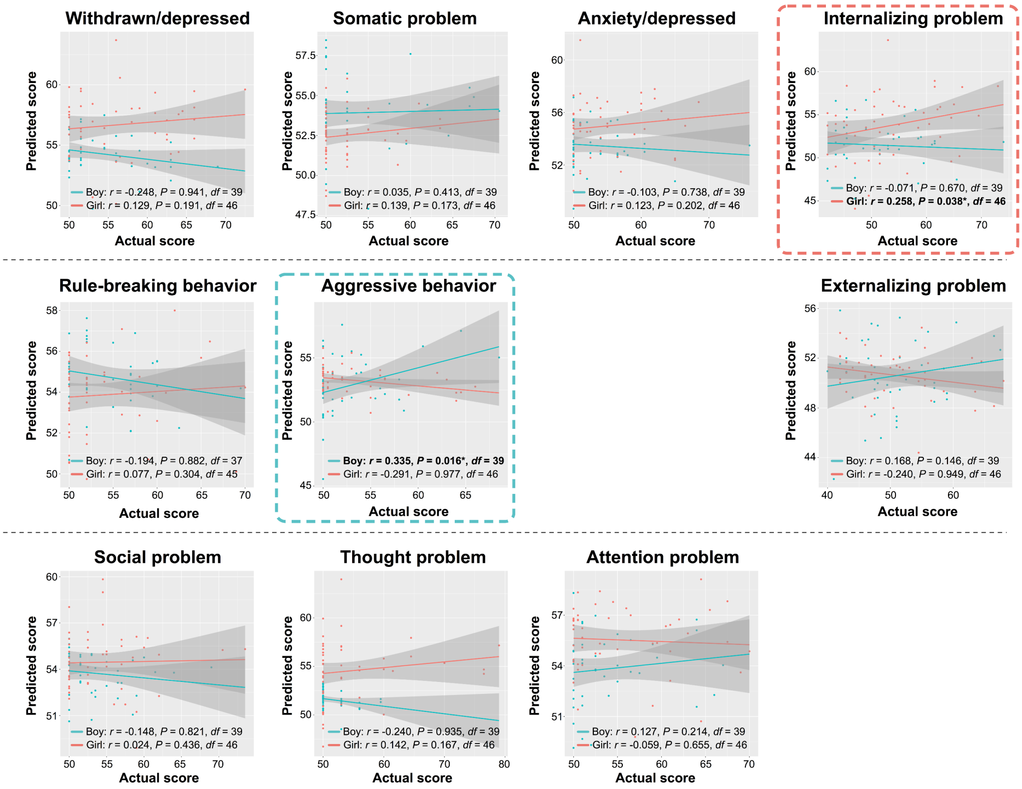


**Fig. S6. Scatter plots of actual versus predicted scores using prediction models with FBCp**
Prediction models for child's psychobehavioral problems were built using independent variables of FBCp. The top, middle, and bottom panels show prediction performances for the CBCL subscales of the internalizing problems, the externalizing problems, and other CBCL subscales (social problem, thought problem, and attention problem), respectively. The pink and green dots represent data points of girl and boy, respectively. Statistical threshold was set to $P < 0.05$. Prediction models exhibited significant prediction accuracies for boy's aggressive behavior [$r = 0.335$, $P = 0.016$, $df = 39$] and girl's internalizing problem [$r = 0.258$, $P = 0.038$, $df = 46$].

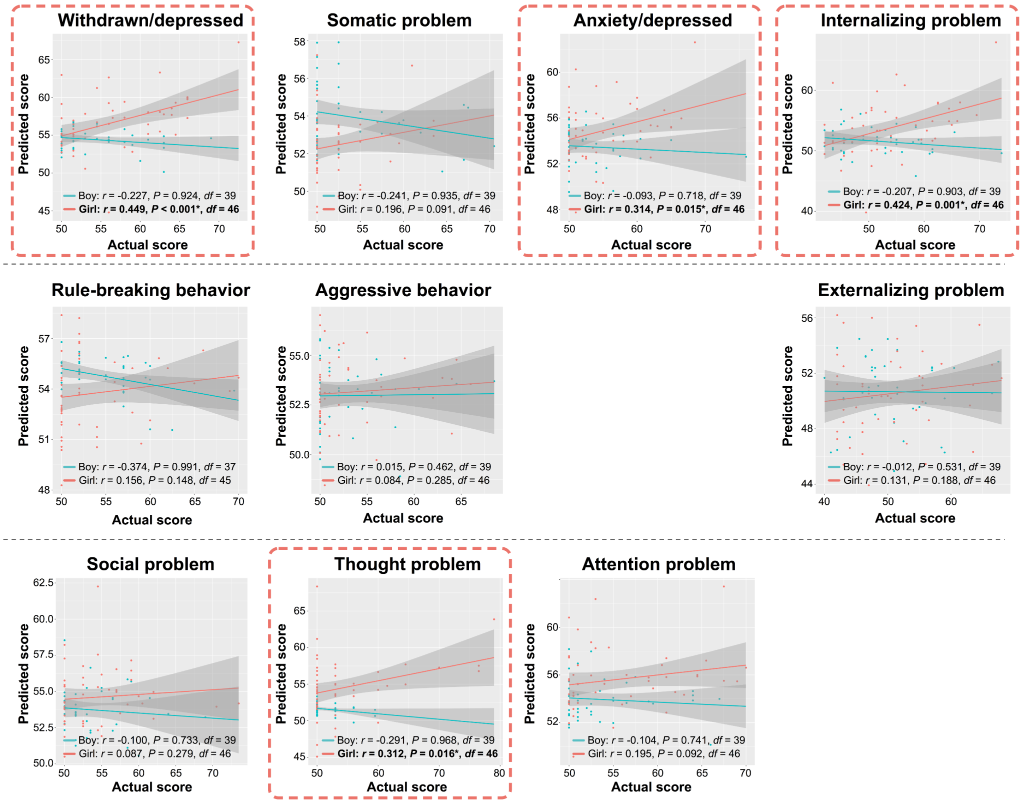


**Fig. S7. Scatter plots of actual versus predicted scores using prediction models with FBCm**
Prediction models for child's psychobehavioral problems were built using independent variables of FBCm. The top, middle, and bottom panels show prediction performances for the CBCL subscales of the internalizing problems, the externalizing problems, and other CBCL subscales (social problem, thought problem, and attention problem), respectively. The pink and green dots denote data points of girl and boy, respectively. Statistical threshold was set to $P < 0.05$. Prediction models exhibited significant prediction accuracies for withdrawn/depressed [$r = 0.449$, $P < 0.001$, $df = 46$], anxiety/depressed [$r = 0.314$, $P = 0.015$, $df = 46$], internalizing problem [$r = 0.424$, $P = 0.001$, $df = 46$], and thought problem [$r = 0.312$, $P = 0.016$, $df = 46$] in girl.

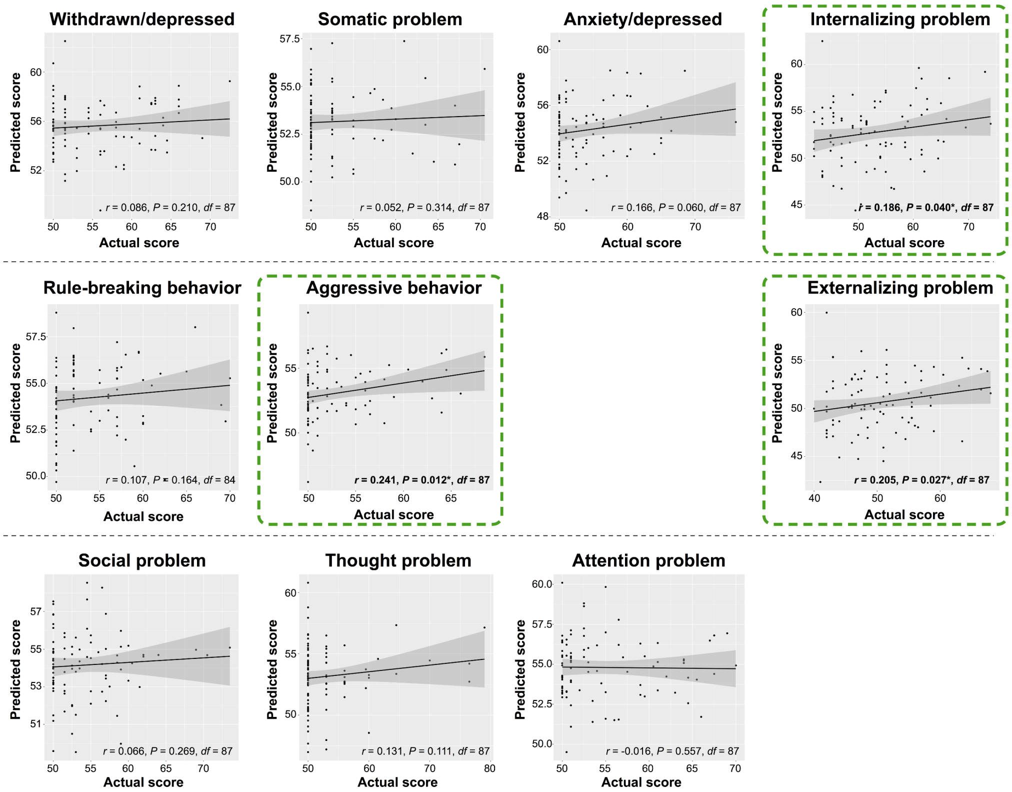


**Fig. S8. Scatter plots of actual versus predicted scores using prediction models that do not account for differences in child's sex**

Prediction models for child's psychobehavioral problems were built using independent variables of both paternal and maternal FBCs without splitting data based on sex. Therefore, a single model was built for each CBCL subscale. The top, middle, and bottom panels show prediction performances for the CBCL subscales of the internalizing problems, the externalizing problems, and other CBCL subscales (social problem, thought problem, and attention problem), respectively. Statistical threshold was set to $P < 0.05$. Prediction models exhibited significant prediction performances for internalizing problem [$r = 0.186$, $P = 0.040$, $df = 87$], aggressive behavior [$r = 0.241$, $P = 0.012$, $df = 87$], and externalizing problem [$r = 0.205$, $P = 0.027$, $df = 87$] but failed to show significant performances for other subscales including withdrawn/depressed, somatic problem, anxiety/depressed, and attention problem.

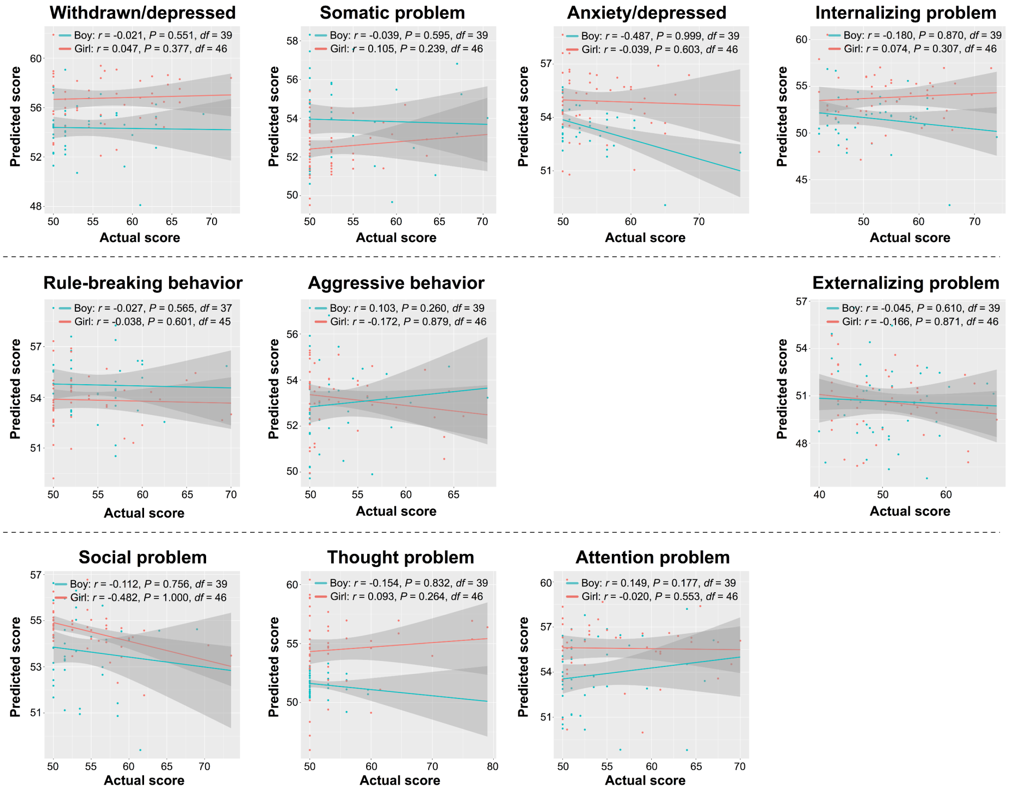


**Fig. S9. Scatter plots of actual versus predicted scores using prediction models that do not account for differences in parental FBCs**

Prediction models for child's psychobehavioral problems were built using FBCs associated with the combined measure of child-parent relationship. The top, middle, and bottom panels show prediction performances for the CBCL subscales of the internalizing problems, the externalizing problems, and other CBCL subscales (social problem, thought problem, and attention problem), respectively. The pink and green dots indicate data points of girl and boy, respectively. Statistical threshold was set to $P < 0.05$. Prediction models could not predict any of the CBCL subscales in boy and girl significantly.

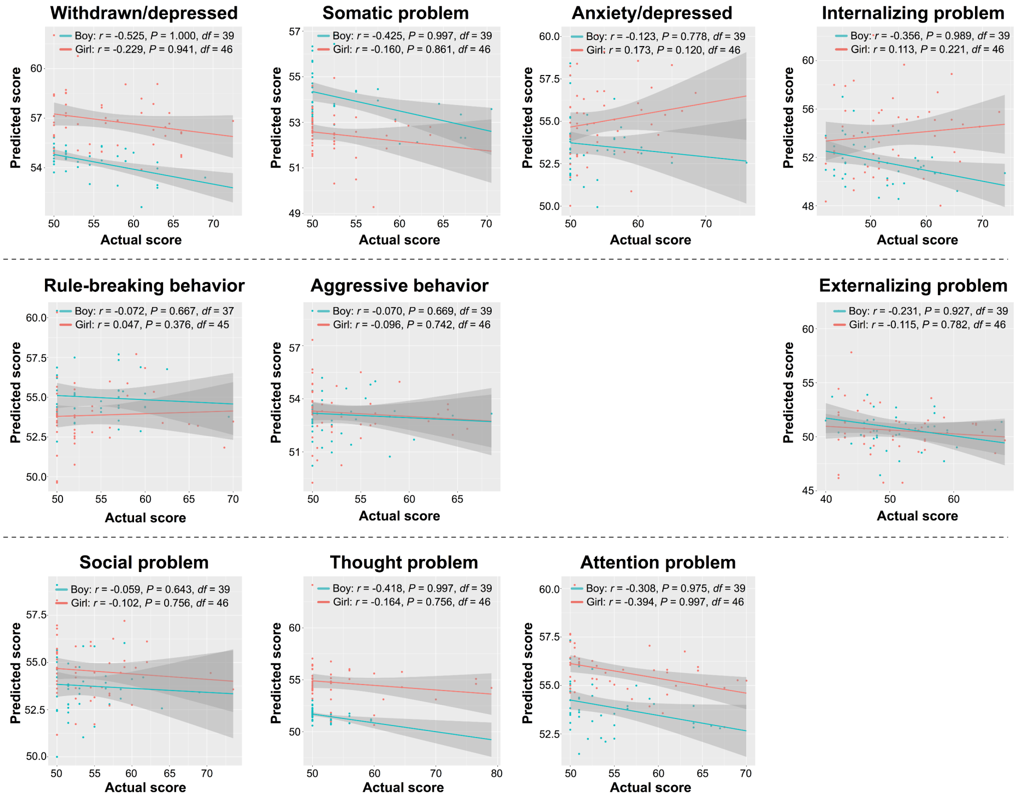


**Fig. S10. Scatter plots of actual versus predicted scores using prediction models with FBCs associated with the peer relationship**

Prediction models for child's psychobehavioral problems were built using FBCs associated with the peer relationship. The top, middle, and bottom panels show prediction performances for the CBCL subscales of the internalizing problems, the externalizing problems, and other CBCL subscales (social problem, thought problem, and attention problem), respectively. The pink and green dots stand for data points of girl and boy, respectively. Statistical threshold was set to $P < 0.05$. Prediction models could not predict any of the CBCL subscales in boy and girl.

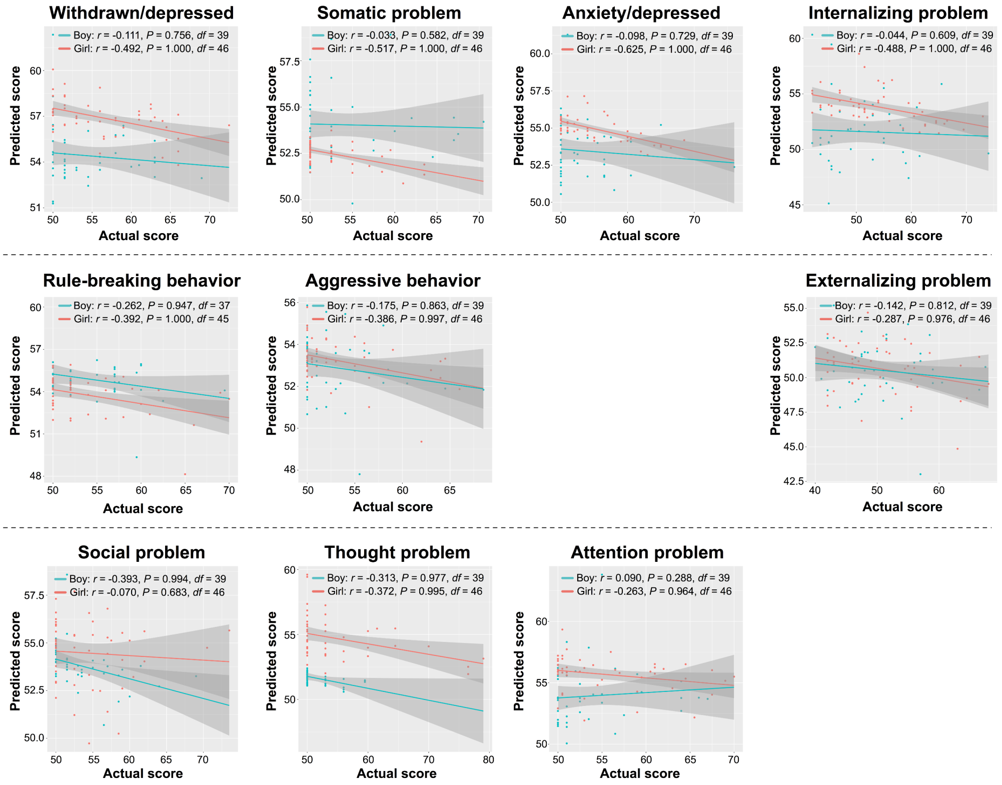


**Fig. S11. Scatter plots of actual versus predicted scores using prediction models with FBCs associated with family SES**

Prediction models for child's psychobehavioral problems were built using FBCs associated with family SES. The top, middle, and bottom panels show prediction performances for the CBCL subscales of the internalizing problems, the externalizing problems, and other CBCL subscales (social problem, thought problem, and attention problem), respectively. The pink and green dots stand for data points of girl and boy, respectively. Statistical threshold was set to $P < 0.05$. Prediction models could not predict any of the CBCL subscales in boy and girl.

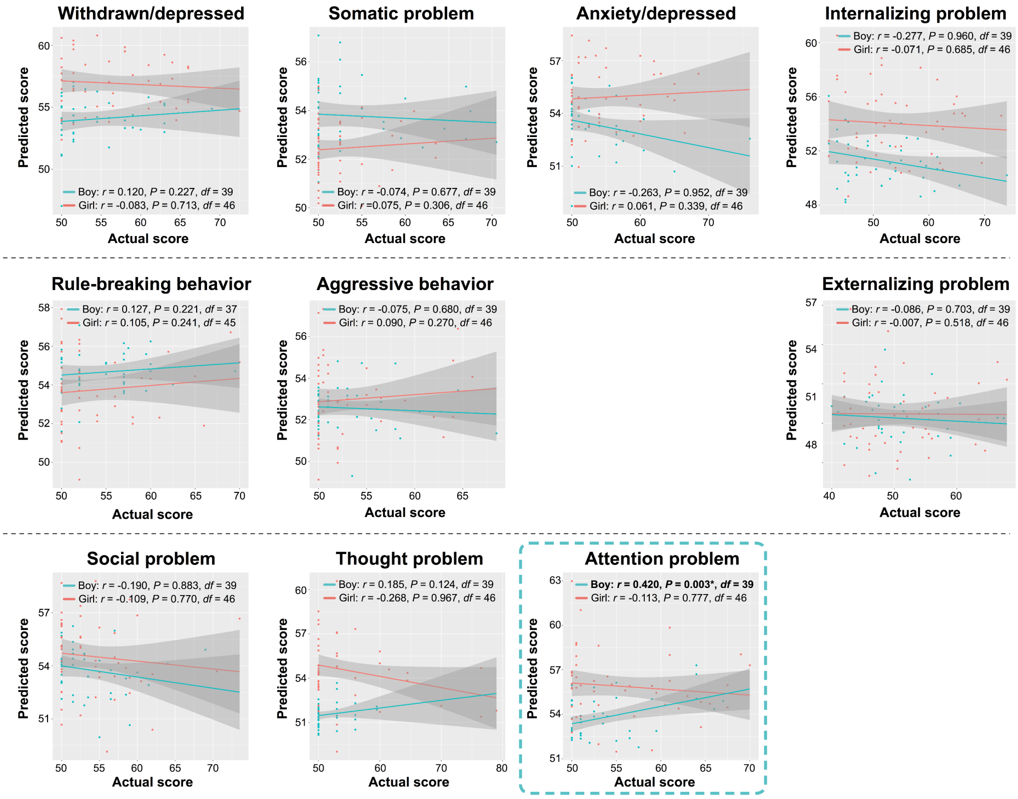


**Fig. S12. Scatter plots of actual versus predicted scores using prediction models with child's psychobehavioral problems themselves**

Prediction models for child's psychobehavioral problems were built using FBCs associated with child's psychobehavioral problems. The top, middle, and bottom panels show prediction performances for the CBCL subscales of the internalizing problems, the externalizing problems, and other CBCL subscales (social problem, thought problem, and attention problem), respectively. The pink and green dots represent girl and boy, respectively. Statistical threshold was set to $P < 0.05$. Prediction models exhibited statistically significant prediction only for boy's attention problem [$r = 0.420$, $P = 0.003$, $df = 39$].

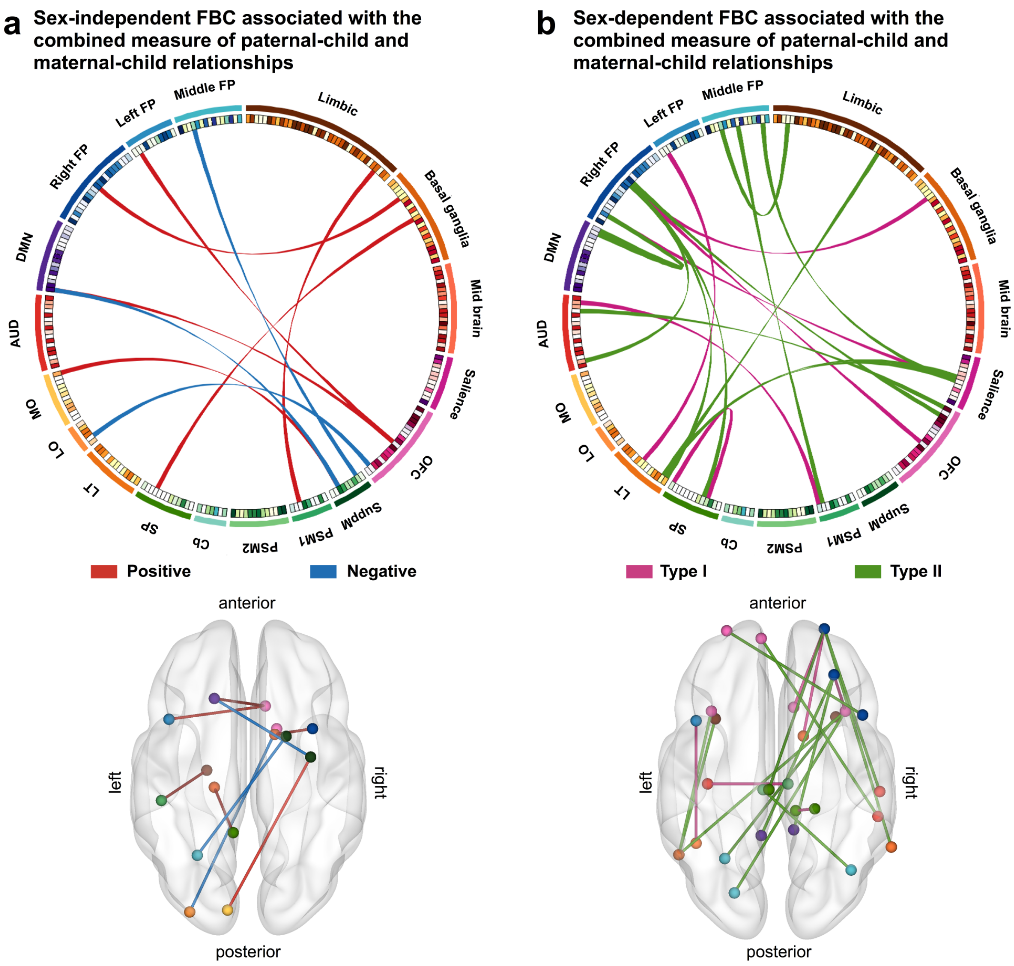


**Fig. S13. Depiction of FBCs associated with the combined measure of the paternal- and maternal-child relationships**

FBCs associated with the combined measure of paternal- and maternal-child relationships were identified by fitting a general linear model. Similar to FBCp or FBCm, the FBC was divided into two major FBCs exhibiting the sex-independent and sex-dependent effects (a, b). Each of the two major FBCs was further subdivided into two sub-FBCs exhibiting positive and negative effects. The red and blue ribbons denote the positive and negative effects of child-parent relationship on FCs in the sex-independent FBC, while the magenta and green ribbons represent these effects in the sex-dependent FBC.

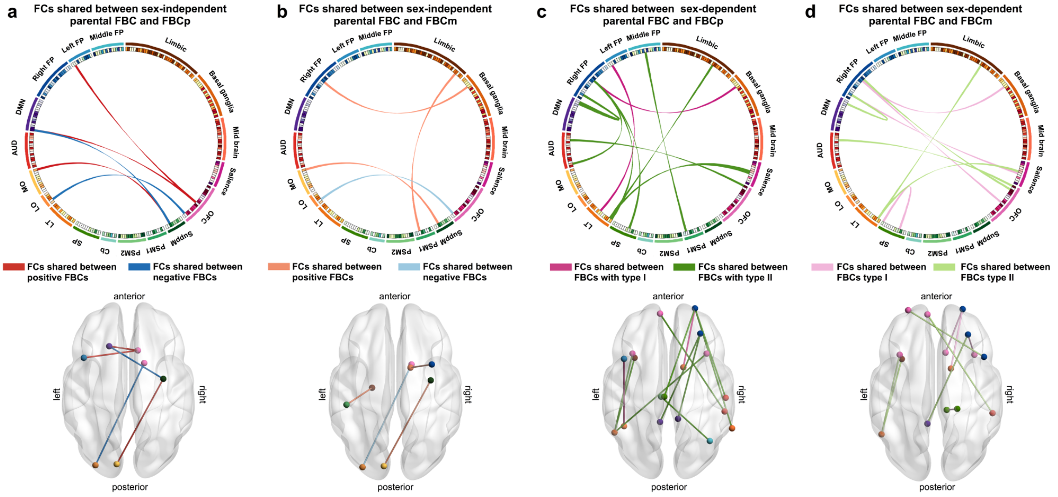


**Fig. S14. Depiction of FCs shared between FBCs associated with the combined measure of paternal- and maternal-child relationships (i.e., 'parental' FBCs) and FBCp and FBCm**
Sex-independent parental FBC shared fives FCs with sex-independent FBCp (a) and four FCs with sex-independent FBCm (b). The red and blue ribbons stand for shared FCs between positive parental FBCs and either positive FBCp or FBCm and those between negative parental FBCs and either negative FBCp or FBCm, respectively. Sex-dependent parental FBC shared 12 FCs with sex-dependent FBCp (c) and nine FCs with sex-dependent FBCm (d). The magenta and blue ribbons indicate shared FCs between parental FBC and either FBCp or FBCm exhibiting the type I effect and those between parental FBC and FBCp or FBCm exhibiting the type II effect, respectively.

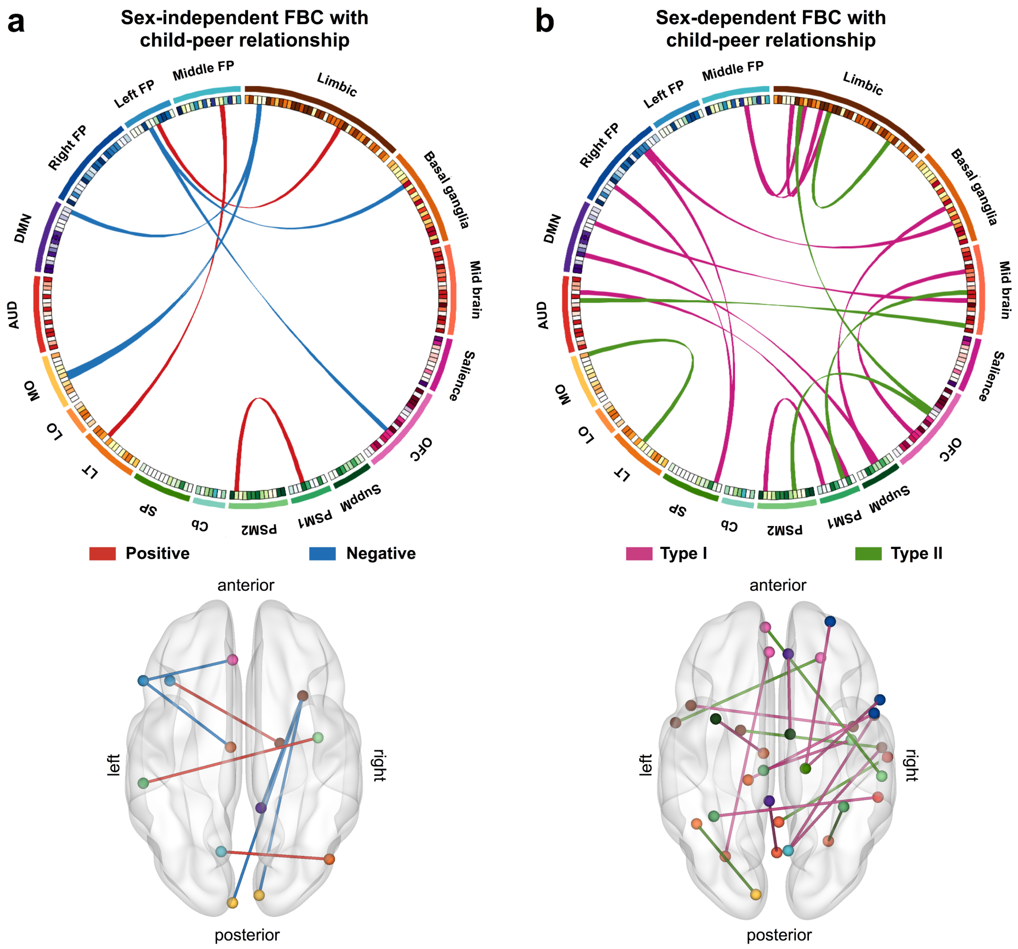


**Fig. S15. Depiction of FBCs associated with the peer relationship**
FBCs associated with the peer relationship were identified by fitting a general linear model in similar to the analyses of the child-parent relationship. FBC associated with the peer relationship was divided into two major FBCs exhibiting the sex-independent and sex-dependent effects (a, b), each of which was further subdivided into two sub-FBCs exhibiting positive and negative effects. The red and blue ribbons stand for the positive and negative effects for sex-independent FBCs, while the magenta and green ribbons denote these effects for sex-dependent FBCs.

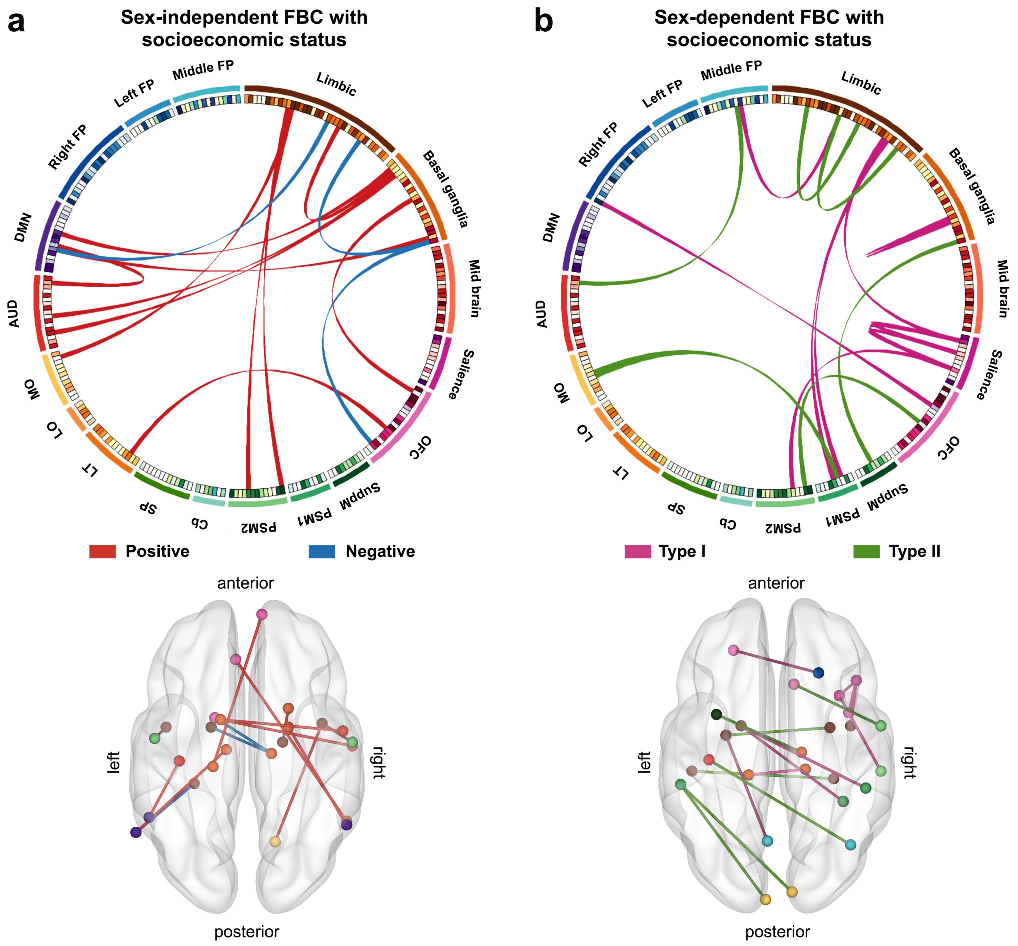


**Fig. S16. Depiction of FBCs associated with family SES**

FBCs associated with family SES were identified by fitting a general linear model in similar to the analyses of the child-parent relationship and of the peer relationship. FBCs associated with family SES were divided into two major FBCs exhibiting the sex-independent and sex-dependent effects (a, b), each of which was further subdivided into two sub-FBCs exhibiting positive and negative effects. The red and blue ribbons indicate the positive and negative effects for sex-independent FBCs, while the magenta and green ribbons stand for these effects for sex-dependent FBCs.

**Table S1. List of FCs involved in FBCs associated with the paternal- and maternal-child relationships.**

| Terminal regions | | | | | | |
|---|---|---|---|---|---|---|
| **Lat.** | **Name** | **Net.** | **Lat.** | **Name** | **Net.** | ***t*-value** |
| **Positive FBCp** | | | | | | |
| L | MFG_L_7_2 | Left FP | R | OrG_R_6_5 | OFC | 3.83 |
| R | PrG_R_6_6 | Right FP | L | LOcC_L_2_1 | MO | 3.77 |
| R | PrG_R_6_2 | SuppM | L | LOcC_L_2_1 | MO | 3.72 |
| L | FuG_L_3_3 | LT | R | LOcC_R_2_2 | Middle FP | 3.68 |
| L | SFG_L_7_2 | DMN | R | OrG_R_6_5 | OFC | 3.67 |
| L | FuG_L_3_3 | LT | L | LOcC_L_2_2 | Middle FP | 3.66 |
| R | SPL_R_5_1 | SP | R | BG_R_6_6 | PSM 2 | 3.44 |
| L | SFG_L_7_2 | DMN | R | Cb_10 | Cerebellum | 3.40 |
| L | FuG_L_3_3 | LT | I | Cb_15 | Midbrain | 3.24 |
| **Negative FBCp** | | | | | | |
| L | SFG_L_7_2 | DMN | R | PrG_R_6_2 | SuppM | -4.71 |
| L | LOcC_L_4_1 | LO | R | BG_R_6_3 | OFC | -4.20 |
| R | PrG_R_6_3 | SuppM | L | PoG_L_4_2 | PSM 2 | -3.77 |
| R | SFG_R_7_4 | SuppM | L | Tha_L_8_6 | BG | -3.74 |
| R | BG_R_6_2 | BG | L | Tha_L_8_6 | BG | -3.71 |
| L | PrG_L_6_1 | PSM 2 | R | STG_R_6_2 | AUD | -3.25 |
| **Positive FBCm** | | | | | | |
| L | PoG_L_4_3 | PSM 1 | L | Hipp_L_2_1 | Limbic | 4.15 |
| L | PhG_L_6_5 | Limbic | R | IPL_R_6_6 | PSM 2 | 3.94 |
| R | MFG_R_7_6 | Right FP | R | BG_R_6_5 | BG | 3.89 |
| L | PhG_L_6_5 | Limbic | L | IPL_L_6_6 | PSM 1 | 3.63 |
| R | PrG_R_6_2 | SuppM | L | LOcC_L_2_1 | MO | 3.61 |
| L | PrG_L_6_4 | SP | R | MVOcC_R_5_2 | MO | 3.58 |
| R | PhG_R_6_3 | Limbic | L | Tha_L_8_3 | BG | 3.49 |
| **Negative FBCm** | | | | | | |

| | | | | | | |
|---|---|---|---|---|---|---|
| L | LOcC_L_4_1 | LO | R | BG_R_6_3 | OFC | -3.55 |
| **FBCp type I** | | | | | | |
| L | IFG_L_6_4 | Left FP | L | IPL_L_6_1 | Middle FP | 3.90 |
| R | MFG_R_7_7 | Right FP | R | BG_R_6_5 | BG | 3.87 |
| L | MFG_L_7_2 | Left FP | L | FuG_L_3_3 | LT | 3.43 |
| **FBCp type II** | | | | | | |
| L | ITG_L_7_2 | LT | L | INS_L_6_2 | Limbic | -5.31 |
| L | ITG_L_7_2 | LT | L | INS_L_6_3 | Salience | -4.44 |
| L | ITG_L_7_2 | LT | R | INS_R_6_3 | Salience | -4.34 |
| R | MFG_R_7_7 | Right FP | R | MTG_R_4_3 | LT | -4.26 |
| L | SFG_L_7_7 | OFC | R | pSTS_R_2_1 | AUD | -4.23 |
| R | MFG_R_7_7 | Right FP | R | STG_R_6_2 | AUD | -4.17 |
| R | IPL_R_6_2 | Middle FP | L | CG_L_7_6 | PSM 1 | -4.13 |
| L | ITG_L_7_2 | LT | R | INS_R_6_2 | Salience | -3.95 |
| R | MFG_R_7_1 | Right FP | L | CG_L_7_4 | DMN | -3.92 |
| R | MFG_R_7_7 | Right FP | L | PCL_L_2_2 | SP | -3.89 |
| R | MFG_R_7_7 | Right FP | L | PoG_L_4_2 | PSM 2 | -3.82 |
| R | MFG_R_7_7 | Right FP | R | PrG_R_6_3 | SuppM | -3.77 |
| L | IFG_L_6_4 | Left FP | L | STG_L_6_5 | Limbic | -3.74 |
| R | MFG_R_7_1 | Right FP | R | CG_R_7_4 | DMN | -3.72 |
| R | IFG_R_6_5 | Salience | L | ITG_L_7_2 | LT | -3.60 |
| R | OrG_R_6_3 | OFC | L | FuG_L_3_3 | LT | -3.55 |
| R | ITG_R_7_5 | LT | R | BG_R_6_1 | OFC | -3.50 |
| R | IPL_R_6_5 | DMN | L | CG_L_7_6 | PSM 1 | -3.45 |
| R | MFG_R_7_7 | Right FP | L | PoG_L_4_4 | SP | -3.36 |
| R | ITG_R_7_5 | LT | L | BG_L_6_3 | OFC | -3.28 |
| **FBCm Type I** | | | | | | |
| R | PCL_R_2_1 | SP | R | PoG_R_4_4 | SP | 4.38 |

| | | | | | | |
|---|---|---|---|---|---|---|
| R | MFG_R_7_7 | Right FP | R | BG_R_6_5 | BG | 4.07 |
| R | MFG_R_7_7 | Right FP | R | OrG_R_6_5 | OFC | 3.87 |
| L | MTG_L_4_1 | AUD | L | LOcC_L_2_2 | Middle FP | 3.45 |
| R | MFG_R_7_1 | Right FP | R | INS_R_6_3 | Salience | 3.41 |
| **FBCm Type II** | | | | | | |
| L | ITG_L_7_2 | LT | L | INS_L_6_2 | Limbic | -5.19 |
| L | MFG_L_7_7 | OFC | R | IFG_R_6_1 | Right FP | -4.75 |
| R | MFG_R_7_1 | Right FP | L | PCun_L_4_4 | DMN | -4.70 |
| L | SFG_L_7_7 | OFC | R | pSTS_R_2_1 | AUD | -4.13 |
| R | MFG_R_7_7 | Right FP | L | CG_L_7_6 | PSM 1 | -4.07 |
| R | MFG_R_7_1 | Right FP | L | CG_L_7_4 | DMN | -3.89 |
| L | ITG_L_7_2 | LT | L | INS_L_6_3 | Salience | -3.84 |
| R | MFG_R_7_1 | Right FP | L | IPL_L_6_2 | Middle FP | -3.55 |
| L | MVOcC_L_5_3 | MO | R | Cb_27 | Midbrain | -3.45 |

The affiliated networks were identified based on the previous meta-analytic study[6]. Statistical threshold was set to $P < 0.05$, false discovery rate correction.

**Abbreviations of networks:** AUD: auditory, BG: basal-ganglia, DMN: default mode network, FP: fronto-parietal, LO: lateral occipital, LT: lateral temporal, MO: medial occipital, OFC: orbitofrontal, PSM: primary sensorimotor, SuppM: supplementary motor, and SP: superior parietal.

**Abbreviations of brain regions:** Amyg: Amygdala, BG: basal ganglia, Cb: cerebellum, CG: cingulate gyrus, Hipp: hippocampus, IFG: inferior frontal gyrus, INS: insular gyrus, IPL: inferior parietal lobule, ITG: inferior temporal gyrus, FuG: fusiform gyrus, LOcC: lateral occipital cortex, MFG: middle frontal gyrus, MTG: middle temporal gyrus, MVOcC: medioventral occipital cortex, OrG: orbital gyrus, PCL: paracentral lobule, Pcun, precuneus, PhG: parahippocampal gyrus, PoG: postcentral gyrus, PrG: precentral gyrus, pSTS: posterior superior temporal sulcus, SFG: superior frontal gyrus, SPL: superior parietal lobule, STG: superior temporal gyrus, and Tha: thalamus.

**Abbreviations of other terms:** L: left, Lat.: laterality, Net.: network, and R: right.

### Table S2. Functional profiles of parental FBCs

| Superordinate domain | FBCp | | | | FBCm | | | |
|---|---|---|---|---|---|---|---|---|
| | Sex-independent effect | | Sex-dependent effect | | Sex-independent effect | | Sex-dependent effect | |
| | Positive | Negative | Type I | Type II | Positive | Negative | Type I | Type II |
| **Action** | 0.029 | 0.028 | 0.011 | 0.032 | 0.029 | 0.004 | 0.009 | 0.024 |
| **Interoception** | 0.007 | 0.006 | 0.004 | 0.011 | 0.009 | 0.002 | 0.005 | 0.007 |
| **Perception** | **0.032** | **0.029** | 0.013 | 0.034 | **0.035** | **0.009** | **0.011** | 0.022 |
| **Cognition** | **0.032** | 0.020 | **0.028** | **0.042** | 0.019 | 0.007 | **0.011** | **0.031** |
| **Emotion** | 0.012 | 0.007 | 0.014 | 0.030 | 0.014 | 0.005 | 0.008 | 0.027 |

Spatial overlap between each FBC and a meta-analytic map for each mental subdomain was calculated and then averaged within each of five superordinate domains.

**Table S3. Detailed functional profiles of parental FBCs**

| | | FBCp | | | | FBCm | | | |
|---|---|---|---|---|---|---|---|---|---|
| | | Sex-independent effect | | Sex-dependent effect | | Sex-independent effect | | Sex-dependent effect | |
| | **Subordinate domains** | **Positive** | **Negative** | **Type I** | **Type II** | **Positive** | **Negative** | **Type I** | **Type II** |
| **Action** | Execution.Speech | 0.030 | **0.042** | 0.016 | 0.047 | 0.035 | 0.007 | 0.007 | 0.018 |
| | Execution | 0.035 | **0.055** | 0.004 | 0.040 | **0.055** | 0.002 | 0.008 | 0.014 |
| | Imagination | 0.032 | **0.038** | 0.012 | 0.013 | **0.046** | 0.000 | 0.003 | 0.011 |
| | Inhibition | 0.033 | 0.018 | 0.019 | 0.051 | 0.023 | 0.003 | **0.024** | **0.053** |
| | Motor_Learning | 0.022 | 0.035 | 0.003 | 0.008 | 0.023 | 0.003 | 0.003 | 0.005 |
| | Observation | 0.038 | 0.022 | 0.026 | 0.051 | 0.022 | 0.017 | 0.006 | 0.027 |
| | Preparation | 0.025 | 0.006 | 0.002 | 0.014 | 0.015 | 0.002 | 0.007 | 0.016 |
| | Rest | 0.014 | 0.005 | 0.010 | 0.034 | 0.011 | 0.002 | 0.013 | **0.047** |
| **Cognition** | Attention | 0.045 | 0.037 | 0.032 | **0.056** | **0.046** | 0.013 | 0.017 | 0.027 |
| | Language.Orthography | 0.042 | 0.023 | **0.046** | 0.045 | 0.008 | **0.022** | 0.013 | 0.036 |
| | Language.Phonology | 0.045 | 0.023 | **0.053** | **0.059** | 0.006 | 0.004 | 0.012 | 0.033 |
| | Language.Semantics | 0.030 | 0.025 | 0.032 | 0.050 | 0.024 | 0.011 | 0.009 | 0.043 |
| | Language.Syntax | 0.020 | 0.010 | 0.029 | 0.036 | 0.007 | 0.000 | 0.015 | 0.036 |
| | Language.Speech | 0.037 | 0.034 | 0.032 | 0.054 | 0.036 | 0.010 | 0.007 | 0.032 |
| | Language | 0.035 | 0.025 | **0.042** | 0.032 | 0.001 | 0.014 | 0.002 | 0.033 |
| | Memory.Explicit | 0.037 | 0.014 | **0.040** | **0.064** | 0.018 | 0.009 | 0.012 | **0.058** |
| | Memory.Implicit | 0.013 | 0.002 | 0.021 | 0.008 | 0.001 | 0.000 | 0.001 | 0.000 |
| | Memory.Working | **0.064** | 0.016 | **0.042** | 0.049 | 0.023 | 0.000 | **0.020** | 0.033 |
| | Memory | 0.017 | 0.006 | 0.022 | 0.022 | 0.035 | 0.005 | 0.001 | 0.014 |
| | Music | 0.018 | **0.047** | 0.001 | 0.046 | 0.027 | 0.003 | 0.005 | 0.020 |
| | Reasoning | 0.041 | 0.024 | 0.035 | 0.054 | 0.021 | **0.017** | **0.023** | 0.042 |
| | Social Cognition | 0.015 | 0.001 | 0.015 | **0.085** | 0.011 | 0.001 | 0.010 | **0.078** |
| | Somatic | 0.012 | 0.020 | 0.010 | 0.017 | 0.029 | 0.003 | 0.006 | 0.010 |
| | Spatial | **0.069** | 0.032 | 0.029 | 0.026 | 0.028 | 0.014 | 0.018 | 0.029 |
| | Temporal | 0.004 | 0.000 | 0.002 | 0.015 | 0.002 | 0.000 | **0.020** | 0.000 |
| **Emotion** | Intensity | 0.002 | 0.002 | 0.007 | 0.013 | 0.001 | 0.000 | 0.004 | 0.006 |
| | Negative.Anger | 0.018 | 0.007 | 0.018 | 0.028 | 0.008 | 0.002 | 0.001 | 0.039 |
| | Negative.Anxiety | 0.006 | 0.001 | 0.014 | 0.046 | 0.024 | 0.002 | 0.018 | 0.035 |
| | Negative.Disgust | 0.015 | 0.013 | 0.009 | 0.045 | 0.018 | 0.008 | 0.008 | 0.043 |
| | Negative.Embarrassment | 0.004 | 0.000 | 0.005 | 0.013 | 0.000 | 0.000 | 0.000 | 0.033 |
| | Negative.Fear | 0.023 | 0.010 | 0.023 | 0.044 | 0.025 | 0.010 | 0.011 | 0.027 |
| | Negative.Guilt | 0.010 | 0.000 | 0.020 | 0.007 | 0.000 | 0.000 | 0.000 | 0.025 |
| | Negative.Punishment-Loss | 0.005 | 0.000 | 0.012 | 0.016 | 0.006 | 0.000 | 0.008 | 0.021 |
| | Negative.Sadness | 0.006 | 0.010 | 0.005 | 0.045 | 0.014 | 0.009 | 0.017 | 0.018 |
| | Negative | 0.002 | 0.008 | 0.004 | 0.027 | 0.023 | 0.015 | 0.001 | 0.032 |
| | Positive.Happiness.Humor | 0.014 | 0.001 | 0.023 | 0.017 | 0.012 | 0.003 | 0.001 | 0.005 |
| | Positive.Happiness | 0.022 | 0.021 | 0.023 | 0.035 | 0.020 | 0.009 | 0.000 | 0.024 |
| | Positive.Reward-Gain | 0.018 | 0.020 | 0.018 | 0.043 | 0.017 | 0.009 | 0.031 | 0.030 |
| | Positive | 0.011 | 0.001 | 0.004 | 0.008 | 0.019 | 0.001 | 0.010 | 0.021 |
| | Valence | 0.004 | 0.001 | 0.008 | 0.014 | 0.012 | 0.000 | 0.002 | 0.013 |
| | Emotion | 0.027 | 0.013 | 0.028 | **0.079** | 0.019 | 0.007 | 0.012 | **0.055** |
| **Interoception** | Baroregulation | 0.002 | 0.005 | 0.001 | 0.006 | 0.004 | 0.000 | 0.004 | 0.013 |
| | Gastrointestinal-Genitourinary_(GI-GU) | 0.005 | 0.006 | 0.000 | 0.022 | 0.017 | 0.000 | 0.011 | 0.007 |
| | Heartbeat Detection | 0.000 | 0.000 | 0.000 | 0.004 | 0.000 | 0.000 | 0.003 | 0.002 |
| | Hunger | 0.012 | 0.002 | 0.020 | 0.016 | 0.012 | 0.001 | 0.000 | 0.009 |
| | Osmoregulation | 0.002 | 0.005 | 0.001 | 0.003 | 0.000 | 0.000 | 0.004 | 0.000 |
| | Respiration Regulation | 0.005 | 0.005 | 0.001 | 0.003 | 0.012 | 0.001 | 0.003 | 0.003 |
| | Sexuality | 0.025 | 0.021 | 0.011 | 0.049 | 0.020 | **0.019** | 0.011 | 0.025 |
| | Sleep | 0.004 | 0.010 | 0.003 | 0.005 | 0.004 | 0.000 | 0.005 | 0.006 |
| | Thermoregulation | 0.000 | 0.001 | 0.000 | 0.000 | 0.003 | 0.000 | 0.001 | 0.002 |
| | Thirst | 0.004 | 0.003 | 0.000 | 0.011 | 0.003 | 0.001 | 0.016 | 0.010 |
| | Vestibular | 0.012 | 0.006 | 0.003 | 0.003 | 0.020 | 0.001 | 0.000 | 0.000 |
| **Perception** | Audition | 0.026 | 0.026 | 0.011 | 0.044 | 0.041 | 0.000 | 0.009 | 0.034 |
| | Gustation | 0.011 | 0.022 | 0.002 | 0.043 | 0.015 | 0.014 | 0.017 | 0.018 |
| | Olfaction | 0.016 | 0.022 | 0.002 | 0.024 | 0.022 | **0.018** | 0.007 | 0.015 |
| | Somesthesis.Pain | 0.020 | 0.036 | 0.002 | 0.042 | **0.049** | 0.003 | 0.007 | 0.016 |
| | Somesthesis | 0.020 | **0.055** | 0.001 | 0.046 | **0.072** | 0.000 | 0.008 | 0.020 |
| | Vision.Color | 0.018 | 0.013 | 0.015 | 0.009 | 0.014 | 0.012 | 0.009 | 0.018 |

| | | | | | | | | |
|---|---|---|---|---|---|---|---|---|
| Vision.Motion | **0.063** | 0.035 | 0.016 | 0.017 | 0.038 | 0.007 | 0.016 | 0.019 |
| Vision.Shape | **0.065** | 0.033 | 0.036 | 0.041 | 0.034 | **0.019** | 0.016 | 0.026 |
| Vision | **0.048** | 0.018 | 0.032 | 0.042 | 0.033 | 0.011 | 0.012 | 0.031 |

Spatial overlap between each FBC and a meta-analytic map for each mental subdomain was calculated.

**Table S4. Summary of statistical properties of input variables used in LiNGAM**

| Scales | Boy | | | Girl | | |
|---|---|---|---|---|---|---|
| | $\gamma$ (95% CI) | $\kappa$ (95% CI) | *P*-value | $\gamma$ (95% CI) | $\kappa$ (95% CI) | *P*-value |
| **The child-parent relationships** | | | | | | |
| NRI score for father | -0.60 (-1.06, -0.07) | -0.29 (-1.13, 1.02) | **<0.001** | -1.08 (-1.55, -0.14) | 1.31 (-0.97, 3.20) | **<0.001** |
| NRI score for mother | - | - | - | -1.58 (-2.12, -0.35) | 2.93 (-1.17, 5.83) | **<0.001** |
| **FBCs associated with the child-parent relationships** | | | | | | |
| Paternal score | -0.25 (-0.71, 0.22) | -1.17 (-1.56, -0.45) | **<0.001** | -0.17 (-0.78, 0.58) | 0.13 (-0.92, 1.02) | **<0.001** |
| Maternal score | - | - | - | -0.24 (-1.11, 0.22) | 1.48 (-0.175, 3.41) | **0.003** |
| **Psychobehavioral problems** | | | | | | |
| Aggressive behavior | 1.85 (0.88, 2.68) | 3.31 (-0.53, 8.23) | **<0.001** | - | - | - |
| Internalizing problem | - | - | - | 0.41 (-0.03, 0.85) | -0.79 (-1.35, 0.15) | **<0.001** |

The variables, γ and *κ*, stand for the skewness and kurtosis of each variable. The 95% confidence interval (CI) was calculated using a bootstrap method with 10,000 iterations implemented in the *R* package "simpleboot".

One-sample Kolmogorov-Smirnov (KS) test was also performed to examine the normality of each variable and P-values were displayed in this table. Statistical threshold was set to $P < 0.05$.

**Table S5. List of FCs involved in FBCs associated with the peer relationship**

| Terminal regions | | | | | | |
|---|---|---|---|---|---|---|
| **Lat.** | **Name** | **Net.** | **Lat.** | **Name** | **Net.** | ***t*-value** |
| **FBC positively associated with the peer relationship** | | | | | | |
| R | IPL_R_6_1 | LT | L | PCun_L_4_3 | Middle FP | 4.24 |
| L | IFG_L_6_5 | Left FP | R | PhG_R_6_4 | Limbic | 3.97 |
| L | IPL_L_6_6 | PSM 1 | R | INS_R_6_5 | PSM 2 | 3.93 |
| **FBC negatively associated with the peer relationship** | | | | | | |
| R | STG_R_6_1 | Limbic | R | MVOcC_R_5_3 | MO | -4.61 |
| R | STG_R_6_1 | Limbic | L | MVOcC_L_5_3 | MO | -3.91 |
| R | STG_R_6_1 | Limbic | R | CG_R_7_4 | DMN | -3.70 |
| L | IFG_L_6_3 | Left FP | L | CG_L_7_3 | OFC | -3.67 |
| L | IFG_L_6_3 | Left FP | L | Tha_L_8_1 | BG | -3.65 |
| **FBC associated with the type I effect of peer relationship** | | | | | | |
| R | MTG_R_4_2 | Limbic | R | PCun_R_4_1 | Middle FP | 4.14 |
| R | ITG_R_7_4 | Limbic | R | PCun_R_4_1 | Middle FP | 4.08 |
| R | pSTS_R_2_1 | AUD | L | SPL_L_5_3 | PSM 1 | 4.01 |
| L | CG_L_7_7 | OFC | L | Cb_14 | Midbrain | 4.01 |
| R | MFG_R_7_3 | Right FP | R | PrG_R_6_4 | SP | 3.89 |
| R | SFG_R_7_5 | SuppM | R | SFG_R_7_6 | DMN | 3.80 |
| R | INS_R_6_5 | PSM 2 | L | CG_L_7_6 | PSM 1 | 3.77 |
| L | CG_L_7_1 | DMN | I | Cb_21 | Midbrain | 3.73 |
| L | STG_L_6_5 | Limbic | R | ITG_R_7_3 | Limbic | 3.70 |
| L | MFG_L_7_6 | SuppM | L | Tha_L_8_4 | BG | 3.57 |
| R | PrG_R_6_6 | Right FP | L | Tha_L_8_6 | BG | 3.47 |
| R | IFG_R_6_6 | Right FP | R | PrG_R_6_4 | SP | 3.37 |

| FBC associated with the type II effect of peer relationship | | | | | | |
|---|---|---|---|---|---|---|
| R | OrG_R_6_3 | OFC | L | MTG_L_4_2 | Limbic | -4.18 |
| L | FuG_L_3_3 | LT | L | LOcC_L_2_1 | MO | -4.04 |
| L | OrG_L_6_4 | OFC | R | IPL_R_6_6 | PSM 2 | -3.77 |
| R | MTG_R_4_4 | AUD | R | Cb_27 | Midbrain | -3.70 |
| R | SPL_R_5_3 | PSM 1 | R | Cb_19 | Midbrain | -3.50 |
| R | ITG_R_7_4 | Limbic | L | Amyg_L_2_1 | Limbic | -3.32 |

The affiliated networks were identified based on the previous meta-analytic study[6]. Statistical threshold was set to $P < 0.05$, false discovery rate correction.

**Abbreviations of networks:** AUD: auditory, BG: basal-ganglia, DMN: default mode network, FP: fronto-parietal, LO: lateral occipital, LT: lateral temporal, MO: medial occipital, OFC: orbitofrontal, PSM: primary sensorimotor, SuppM: supplementary motor, and SP: superior parietal.

**Abbreviations of brain regions:** Amyg: Amygdala, BG: basal ganglia, Cb: cerebellum, CG: cingulate gyrus, Hipp: hippocampus, IFG: inferior frontal gyrus, INS: insular gyrus, IPL: inferior parietal lobule, ITG: inferior temporal gyrus, FuG: fusiform gyrus, LOcC: lateral occipital cortex, MFG: middle frontal gyrus, MTG: middle temporal gyrus, MVOcC: medioventral occipital cortex, OrG: orbital gyrus, PCL: paracentral lobule, Pcun, precuneus, PhG: parahippocampal gyrus, PoG: postcentral gyrus, PrG: precentral gyrus, pSTS: posterior superior temporal sulcus, SFG: superior frontal gyrus, SPL: superior parietal lobule, STG: superior temporal gyrus, and Tha: thalamus.

**Abbreviations of other terms:** L: left, Lat.: laterality, Net.: network, and R: right.

**Table S6. List of FCs involved in FBCs associated with family SES**

| Terminal regions | | | | | | |
|---|---|---|---|---|---|---|
| **Lat.** | **Name** | **Net.** | **Lat.** | **Name** | **Net.** | ***t*-value** |
| **FBC positively associated with the SES** | | | | | | |
| L | MTG_L_4_3 | DMN | L | Tha_L_8_8 | BG | 4.72 |
| R | ITG_R_7_2 | LT | L | CG_L_7_3 | OFC | 4.07 |
| R | PhG_R_6_4 | Limbic | R | BG_R_6_4 | BG | 3.99 |
| R | ITG_R_7_3 | Limbic | R | MVOcC_R_5_5 | MO | 3.96 |
| R | STG_R_6_3 | AUD | L | BG_L_6_5 | BG | 3.89 |
| L | PrG_L_6_1 | PSM 2 | L | ITG_L_7_3 | Limbic | 3.87 |
| R | ITG_R_7_3 | Limbic | R | PoG_R_4_2 | PSM 2 | 3.76 |
| L | pSTS_L_2_2 | DMN | L | INS_L_6_1 | AUD | 3.69 |
| R | IPL_R_6_5 | DMN | R | BG_R_6_2 | BG | 3.65 |
| R | SFG_R_7_7 | OFC | L | Tha_L_8_3 | BG | 3.61 |
| R | STG_R_6_6 | AUD | L | BG_L_6_5 | BG | 3.21 |
| **FBC negatively associated with the SES** | | | | | | |
| L | BG_L_6_3 | OFC | R | Tha_R_8_8 | BG | -4.01 |
| L | Amyg_L_2_1 | Limbic | R | Tha_R_8_8 | BG | -3.91 |
| L | MTG_L_4_3 | DMN | L | PhG_L_6_3 | Limbic | -3.79 |
| **FBC associated with the type I effect of SES** | | | | | | |
| R | IPL_R_6_3 | PSM 1 | L | Amyg_L_2_1 | Limbic | 4.41 |
| R | INS_R_6_2 | Salience | R | INS_R_6_6 | Salience | 4.36 |
| L | PhG_L_6_1 | Limbic | L | PCun_L_4_1 | Middle FP | 4.21 |
| R | SPL_R_5_3 | PSM 1 | L | Amyg_L_2_1 | Limbic | 3.84 |
| R | IPL_R_6_6 | PSM 2 | R | INS_R_6_6 | Salience | 3.71 |
| R | Tha_R_8_5 | BG | L | Tha_L_8_6 | BG | 3.66 |
| R | SFG_R_7_2 | Right FP | L | OrG_L_6_3 | OFC | 3.55 |

| | | | | | | |
|---|---|---|---|---|---|---|
| R | IFG_R_6_5 | Salience | R | INS_R_6_2 | Salience | 3.50 |
| R | IFG_R_6_5 | Salience | R | INS_R_6_6 | Salience | 3.50 |
| R | IFG_R_6_5 | Salience | R | INS_R_6_4 | Limbic | 3.39 |
| **FBC associated with the type II effect of SES** | | | | | | |
| L | PhG_L_6_1 | Limbic | R | Amyg_R_2_2 | Limbic | -4.35 |
| L | MFG_L_7_6 | SuppM | R | Tha_R_8_8 | BG | -4.30 |
| R | IPL_R_6_2 | Middle FP | L | INS_L_6_1 | AUD | -3.99 |
| L | IPL_L_6_3 | PSM 1 | R | MVOcC_R_5_3 | MO | -3.93 |
| L | ITG_L_7_1 | Limbic | R | PhG_R_6_3 | Limbic | -3.77 |
| L | IPL_L_6_3 | PSM 1 | L | MVOcC_L_5_3 | MO | -3.48 |
| R | OrG_R_6_5 | OFC | R | PrG_R_6_1 | PSM 2 | -3.47 |

The affiliated networks were identified based on the previous meta-analytic study[6]. Statistical threshold was set to $P < 0.05$, false discovery rate correction.

**Abbreviations of networks:** AUD: auditory, BG: basal-ganglia, DMN: default mode network, FP: fronto-parietal, LO: lateral occipital, LT: lateral temporal, MO: medial occipital, OFC: orbitofrontal, PSM: primary sensorimotor, SuppM: supplementary motor, and SP: superior parietal.

**Abbreviations of brain regions:** Amyg: Amygdala, BG: basal ganglia, Cb: cerebellum, CG: cingulate gyrus, Hipp: hippocampus, IFG: inferior frontal gyrus, INS: insular gyrus, IPL: inferior parietal lobule, ITG: inferior temporal gyrus, FuG: fusiform gyrus, LOcC: lateral occipital cortex, MFG: middle frontal gyrus, MTG: middle temporal gyrus, MVOcC: medioventral occipital cortex, OrG: orbital gyrus, PCL: paracentral lobule, Pcun, precuneus, PhG: parahippocampal gyrus, PoG: postcentral gyrus, PrG: precentral gyrus, pSTS: posterior superior temporal sulcus, SFG: superior frontal gyrus, SPL: superior parietal lobule, STG: superior temporal gyrus, and Tha: thalamus.

**Abbreviations of other terms:** L: left, Lat.: laterality, Net.: network, and R: right.
